# Serial depletion of Baltic herring since the Viking Age revealed by ancient DNA and population genomics

**DOI:** 10.1101/2022.05.18.492457

**Authors:** Lane M. Atmore, Lourdes Martínez-García, Daniel Makowiecki, Carl André, Lembi Lõugas, James H. Barrett, Bastiaan Star

## Abstract

Marine resource consumption has been a key component in European diet and culture since the Middle Ages, when fish consumption increased dramatically. Yet, the early origins of marine industries and the long-term ecological consequences of historical and contemporary fisheries remain debated. The Baltic Sea was home to the first “industrial” fishery ∼800 years ago targeting the Baltic herring, a species that is still economically and culturally important today. We combine modern whole genome data with ancient DNA (aDNA) to identify the first known long-distance herring trade in the region, illustrating that large-scale fish trade began during the Viking Age. We resolve population structure within the Baltic and observe demographic independence for four local herring stocks over at least 200 generations. It has been suggested that overfishing at Øresund in the 16^th^ century resulted in a demographic shift from autumn-spawning to spring-spawning herring dominance in the Baltic. We show that while the Øresund fishery had a negative impact on the western Baltic herring stock, the existence of autumn-spawning refugia in other regions of the Baltic delayed the demographic shift to spring spawning dominance until the 20^th^ century. Importantly, modelling demographic trajectories over time, we identify a consistent pattern of serial depletion within the Baltic that is associated with changes in fishing pressure and climate, and conclude that herring exploitation at both historical and recent intensities is not sustainable. Our results highlight the complex and enduring impacts humans have had on the marine environment well before the industrial era.

## Introduction

During the Middle Ages, diets throughout Europe shifted to rely more heavily on marine fish^1^. Increased demand for fish protein followed the rise of urbanism and the spread of Christian fasting practices in Europe^1–3^. Herring (*Clupea harengus*), was a particularly valued species at the time due to its ability to be preserved and sold in quantity at market, its massive spawning aggregations that provided easy capture, and its high fat content^4,5^. Long-distance herring trade, however, was limited by economical access to salt, which is required to keep fresh-caught herring from spoiling^6^. One of the earliest fisheries that had access to both salt and coastal herring spawning aggregations occurred in the western Baltic. The Baltic Sea has long been under intense exploitation pressure^7^ and is particularly vulnerable to both climate change^8^ and intensified extraction^9^. It therefore provides an excellent opportunity to evaluate shifting impacts of human exploitation and changing climate on marine species^10^. According to historical data, the earliest commercial herring fishing operations in the Baltic took place on the island of Rügen in the early 12^th^ century^11^, an operation that was succeeded by the first fishery in Europe to reach the level of a true industry: the Øresund fishery^7,12,13,13^.

The Øresund fishery (here broadly defined) operated between Denmark and southern Sweden, corresponding to ICES subdivisions 22-24, with most activity concentrated in 23-24^6,15^ (see Fig 1A). During the peak of the Øresund fishery, herring were traded across Europe, reaching as far away as Italy and northern Norway^16^. Yet, the nature, origin, and timing of onset of this commercial fishery remains unclear^7,17^. The famous account of the Anglo-Saxon Wulfstan^18^ reports a voyage c.880 CE between the trading ports Hedeby (in modern Schleswig-Holstein, Germany) and Truso (identified at the archaeological site of Janów Pomorski) near Elbląg in Poland. Zooarchaeological evidence from a herring bone assemblage dated c.800-850 CE from Janów Pomorski might be indicative of such early medieval fish trade in the same direction due to the absence of cleithra – a bone element often removed during medieval commercial processing^17,19^. This is a hypothesis that can now be tested using ancient DNA (aDNA).

**Figure 1.**
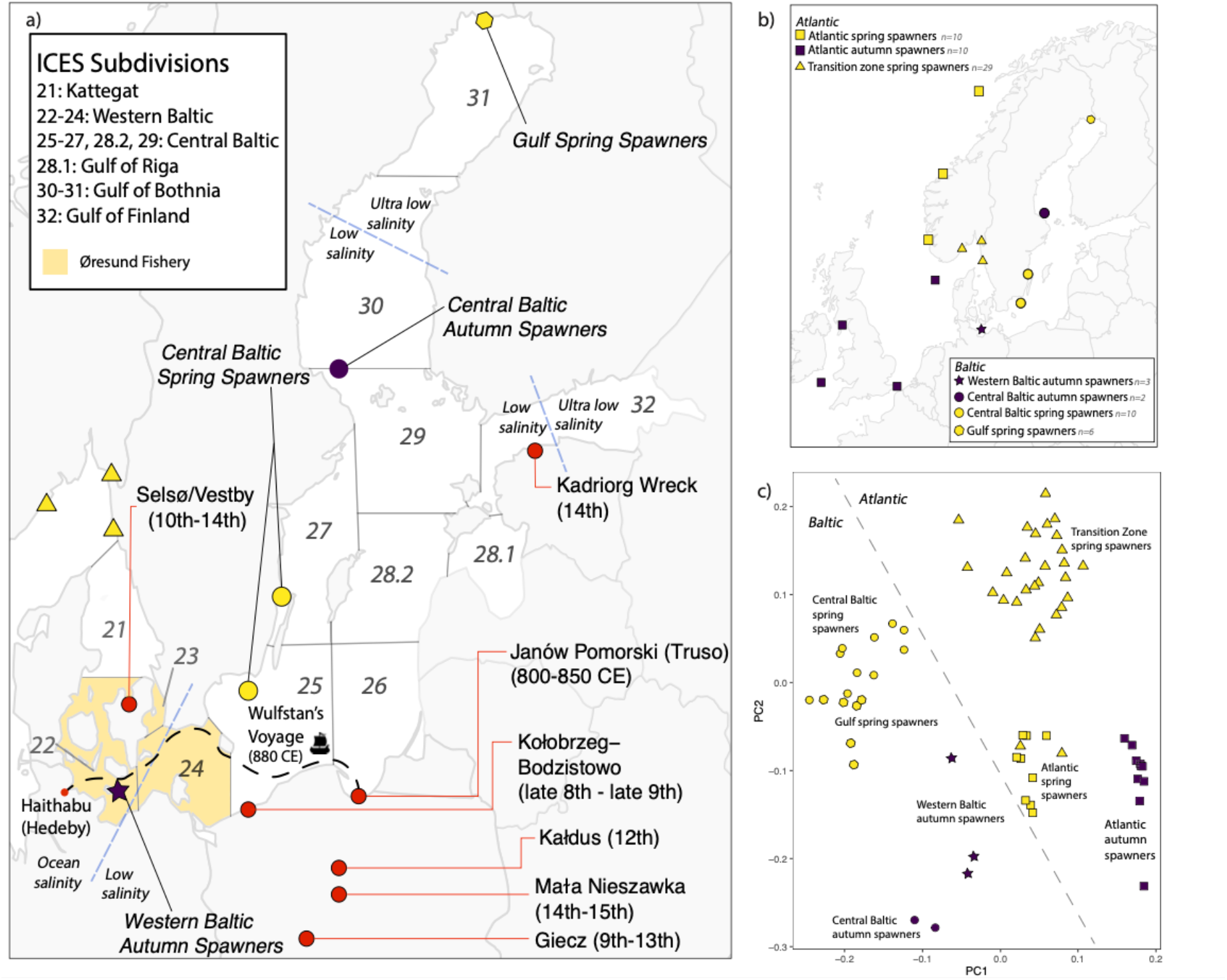
Archaeological sites, sample location, and population structure of Baltic herring. a) Baltic Sea with ICES management subdivisions indicated. Named regions corresponding to ICES subdivisions are noted in the figure legend. Spring and autumn spawning herring occur in all subdivisions. Archaeological sites and their dates are noted as well as the sampling locations for modern genome data. The approximate area targeted by the Øresund fishery (broadly defined) in the Middle Ages is highlighted in yellow (ICES 22-24), although it concentrated in subdivisions 23-24. Approximate locations of salinity boundaries between high (ocean), low, and ultra-low salinity are marked by blue dashed lines. Wulfstan’s voyage from Hedeby to Truso c.880 CE is noted in black. b) Sample locations of herring specimens used for this study. These specimens are sourced from publicly available material (n=22) and include newly-generated genomes (n=53). Color denotes spawning season, indicating near-complete coverage of the major herring populations identified by Han et al.^33^; c) PCA of the entire modern nuclear genome using 10,368,446 SNPs. The modern genome shows segregation on PC1 based on Atlantic vs Baltic and on PC1 and spawning season on PC2. The Atlantic spring spawners are divided into two groups, with the transition zone between the Baltic and Atlantic separating on PC1. This is likely due to the presence of Norwegian fjord herring in the transition zone cluster, which show signatures of adaptation similar to the Baltic herring due to the brackish fjord conditions^88^. Winter and autumn spawning specimens from the Atlantic cluster together. Gulf spring spawners and central Baltic spring spawners show substructure within the Baltic spring spawning metapopulation and western Baltic autumn spawners cluster away from the central Baltic autumn spawners.

Under the control of Denmark and the Hanseatic League, the Øresund fishery was most active during the 13^th^-16^th^ centuries. Exact quantities of catch are still under debate^6,7^, but the Øresund fishery likely surpassed contemporary western Baltic fisheries, with catches estimated at 50 000 tonnes per annum at its peak^6,7,12^. In contrast, early 20^th^ century landings from the Øresund were between 100 and 10 000 t^20^ and more recent landings have been in the order of 10 000 – 20 000 t^15^. Unlike current Baltic fisheries, the Øresund fishery focused exclusively on autumn-spawning herring, with a restricted fishing season between August-October to target the spawning aggregation^6,21^. Today there are autumn spawners extant in the Baltic, but none of these populations are large enough to support a commercial industry^15^. Nowadays, Baltic herring fisheries instead target the smaller-bodied spring-spawners, which are ecologically distinct from autumn-spawners^22^. Although both autumn- and spring-spawning populations co-exist whereever herring are found, in each area only one of these distinct ecotypes appears to dominate numerically (e.g. spring spawners in the Norwegian Sea and autumn spawners in the North Sea)^23,24^. This alternating pattern of dominance in today’s oceans, and the known historical shift towards targeting spring-instead of autumn-spawning stocks, indicate that a dramatic demographic and ecological shift has occurred in the Baltic ecosystem. Nonetheless the timing and nature of this shift is currently poorly characterized.

The Øresund fishery collapsed suddenly between 1570 and 1580. Catches in this decade fell by over 90% from > 60 000 t to < 5 000 t per annum, with only marginal operations continuing into the 17^th^ century^6^. After this, commercial fishing operations ceased in the area until the 20^th^ century^6,25^. European herring production moved to the North Sea^26^, a transition that is generally linked to the decline in the Øresund fishery landings. Yet, whether this decline was due to ecological changes or driven by market and political factors is still under debate^27^. It has recently been argued that the impetus for the collapse of the Øresund fishery was the disappearance of the western Baltic autumn-spawning herring due to overfishing^6^. Based on historical catch records, it has been suggested that landings per annum over the course of the Øresund fishery pushed the stock past modern maximum-sustainable yield (MSY) in the 16^th^ century^5,15^, resulting in stock collapse. It has been proposed that such collapse may have led to a demographic dominance of spring-spawning herring as early as the 16^th^ century^6^. Yet, catch records indicate that autumn-spawners constituted up to 90% of Baltic herring landings as late as 1927^28^, and autumn-spawning fisheries were supported in the gulfs of Finland, Riga, and Bothnia into the 1970s^29^. Past research regarding herring stock collapses has shown that the species is vulnerable to overfishing despite its natural abundance^25,29,29^; the medieval Øresund fishery could well have had a real ecological impact on the herring stock. However, whether this fishery caused a Baltic-wide demographic shift, a short-term local extirpation, or more lasting impacts remains unclear. The impacts of fisheries that succeeded the medieval Øresund industry are also uncharted. Moreover, changing climate in the Baltic during periods such as the Medieval Climate Anomaly and the Little Ice Age likely contributed to herring population dynamics as well^30,31^. Therefore, a holistic evaluation of complex herring ecology, human impacts and the dynamic Baltic ecosystem is merited.

The Baltic exhibits a stark salinity gradient from near-ocean levels where it connects with the Kattegat, to near-freshwater in the gulfs^32^. Baltic herring is a schooling fish with enormous population sizes, with current stock estimates in the order of millions of tonnes of spawning stock biomass^15^. Herring show strong genetic adaptation for spawning season and adaptation to differing salinity conditions^22,33–36^, ultimately segregating into two distinct metapopulations: spring spawners and autumn spawners^37,38^. Spring spawners are smaller and mature more quickly than autumn spawners, and prefer to spawn in coastal areas as opposed to the deeper waters used by the slow-growing, larger autumn spawners^29,39^. Given the larger size of autumn-spawning herring, we expect that earlier fishing efforts prioritized these populations over the smaller spring-spawning aggregations. Each metapopulation is further separated into management stocks living in the western Baltic (ICES subdivisions 21-24), central Baltic (ICES subdivisions 25-27, 28.1, 29), and gulfs (ICES subdivisions 28.1, 30-32) (see Fig 1A)^32,29,40^. The western Baltic autumn-spawning stock is the population targeted by the Øresund fishery. Western Baltic herring (both spring and autumn spawners) spawn in coastal areas of the Kattegat and southwestern Baltic^30,41^, and are likely reproductively isolated populations from the rest of the Baltic^42,43^.

Herring remain an important cultural and commercial industry in the Baltic^15^. As Europe attempts to shift towards a more “sustainable” diet based on aquaculture and marine resources, herring are becoming a key species in the proposed fight to reduce global carbon footprints^44,45,46,47,48,49^. Baltic herring are facing the brunt of our attempts to reduce land-based meat consumption both through a regional increase in fishmeal production for aquaculture^49^ as well as direct consumption. In the European Union, there have been calls to increase understanding of ocean ecosystems to assess the viability of feeding the world from the ocean^47^. Ecologists have already noted the disconnect between the reality of modern ocean environments and governments’ proposed “sustainable” new marine industries^48^, although it is unclear whether this analysis holds true in all ecosystems or over time.

Here we publish the first long time series of genome-wide ancient DNA data for Baltic herring, analyzing 40 archaeological herring specimens from seven sites in Poland, Denmark, and Estonia that span the period from 750-1600 CE. Using *BAMscorer*, software specifically designed for population assignment using ultra-low-coverage sequence data^50^, we identify the biological origin of each specimen, illustrating the change in fishery targets throughout time, and evaluating the onset of long-range fish trade in the Baltic. We use modern whole-genome resequencing data – including 53 new sequences generated for this study– to investigate population structure and model recent past demography for each of the Baltic herring populations in our data (see Fig 1C). We assess the difference in western Baltic autumn-spawning herring outcomes for the Øresund fishery in comparison with more recent fishing efforts and provide insight into the timing of herring population turn-over in the Baltic. We compare the demographic trajectories of four Baltic herring populations to known historical events, including changed fishing practices and temporal changes in sea surface temperature, to assess the impact of these events on herring population size and the long-term sustainability of this iconic industry.

## Results

### Population Structure

We resolve modern population structure in a PCA based on 10 million genome-wide SNPs in 68 modern individuals (Fig 1A,B), consistently separating population based on geography (Baltic or Atlantic) and spawning season (Fig 1C). We further observe fine-scaled population structure within spawning ecotypes in the Baltic, with Central Baltic and Gulf spring spawners, as well as Central Baltic and Western Baltic autumn spawners clustering closer to their respective sample locations (Fig 1C). This fine-scaled structure is also supported by higher levels of relatedness within each sample location and subsequently lowered levels of relatedness in these subpopulations when grouped as metapopulations (see Supplement S1, Fig S10). The modern mitogenomes revealed three major clades (IQ-TREE^51,52^) that exhibited no association with geography or environment across the Baltic and Atlantic, in accordance with previously-published results^43^. All archaeological mitogenomes clustered with the modern Atlantic and Baltic samples, identifying these as herring (Fig S12). Genetic diversity (*π*, VCFtools v0.1.16^53^) across the genome was relatively high in both metapopulations, and significantly higher for autumn-spawning herring (Fig S13). This higher diversity indicates a larger effective population size over time for the autumn spawners relative to spring spawners.

### Population Assignment

All 40 archaeological samples were classified using *BAMscorer* as autumn-spawning herring, with nearly all exhibiting the chromosome 12 inversion haplotype (BB) associated with Baltic herring (Fig 2). Spawning season and chromosome 12 inversion type could be assigned with as few as 50,000 reads, while salinity adaptation required 60,000 reads for accurate assignment (Figs S7-S9). Two specimens from Truso had too few reads to properly assign a salinity adaptation but could be assigned for chromosome 12 inversion type and spawning season (Fig 2, Supplementary Dataset S2). Salinity adaptation was mixed between all sites, with earlier sites showing some individuals with higher salinity adaptation, which is indicative of populations that spend part of their annual cycle in the Skagerrak and North Sea, such as the western Baltic autumn spawning herring, and later sites showing more low-salinity-adapted individuals (Fig 2, Supplementary Dataset S2). Indeed, the oldest samples in the dataset – from Truso in Poland, dating to 800-850 CE^19^ – showed nearly half of samples stemming from high-salinity adapted populations. In contrast, the sites dating to later periods were more dominated by low-salinity adapted autumn spawners. This change indicates a spatial shift in target population for Baltic fisheries over time from western to central Baltic, and points to the existence of refugia for the autumn-spawning herring metapopulation outside of the Øresund.

**Figure 2.**
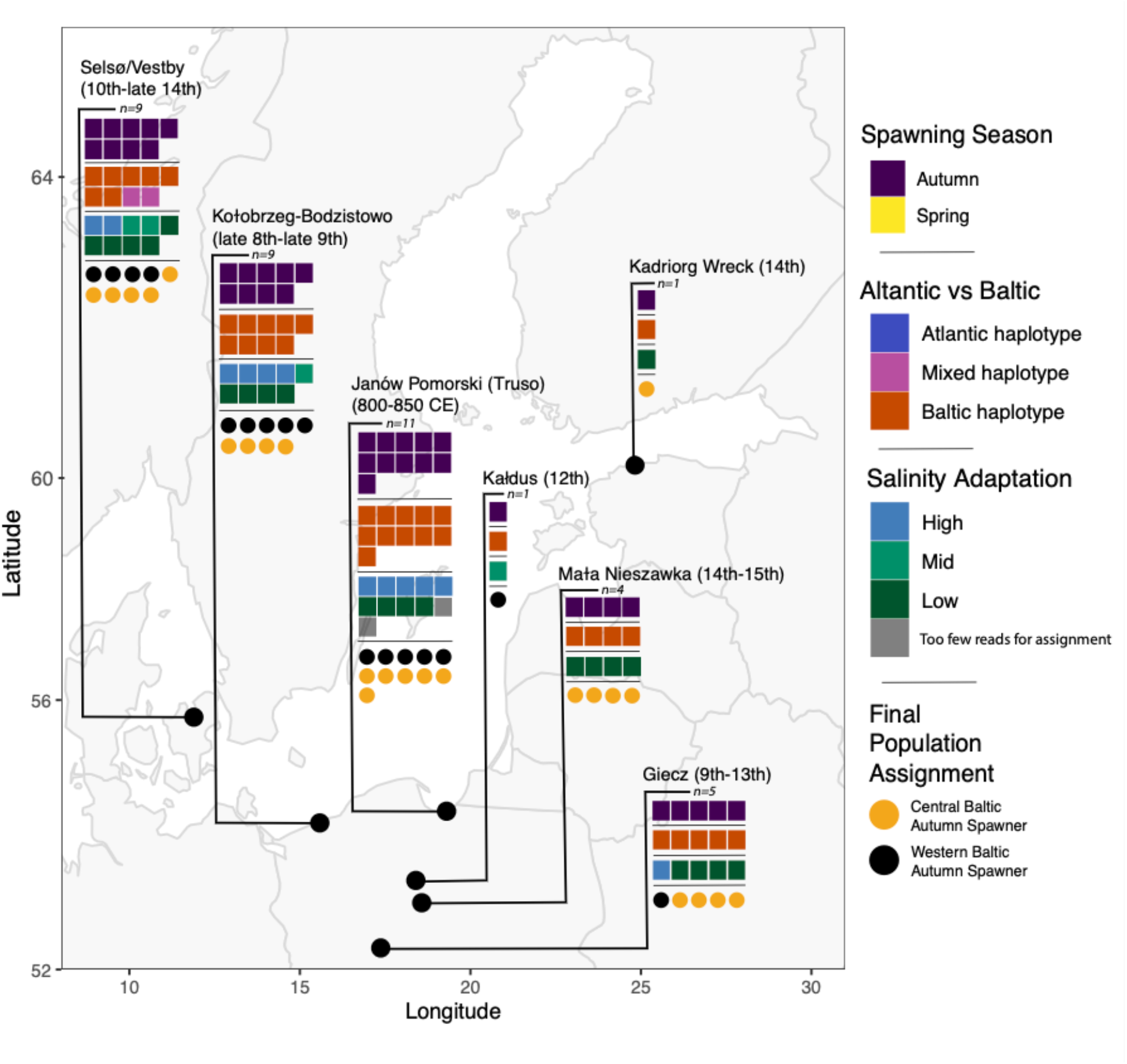
Population assignment results on archaeological herring specimens. Each individual is represented by a single square for each assignment test. The three assignment tests (spawning season, chromosome 12 inversion type, salinity adaptation) are then combined to make a final population assignment illustrated by the row of circles for each site. Western Baltic specimens are present in the earlier sites from around Poland whereas later sites and contexts trend towards central Baltic specimens. Context-specific results can be found in Supplementary dataset S1.

### Demographic Analysis

Both Baltic autumn-spawning populations showed fewer and shorter runs of homozygosity across all time bins (Fig S14). This pattern indicates either a large effective population size or population growth^54^. The Baltic spring spawning populations exhibited signs of smaller effective population size and/or population decline across all time bins. No ROH >330KB were found in any population, likely due to the high recombination rate in herring (2.54 CM/Mb)^35^ and small sample size for this analysis. The combined ROH for all time periods indicates a large effective population size in the past for autumn-spawning herring and smaller effective population size over time for spring-spawning herring.

Highly divergent demographic trajectories are obtained using *gone*^55^ for each of the four Baltic populations under study (Fig 3): Central Baltic Spring Spawners (CBSS), Central Baltic Autumn Spawners (CBAS), Western Baltic Autumn Spawners (WBAS), and Gulf Spring Spawners (GSS). These results illustrate long-term demographic independence of the four main herring stocks in the Baltic. The WBAS appear to have begun declining ∼800YBP, which corresponds to the onset of the Øresund fishery (Fig 3). The WBAS population never fully recovered from this decline, gradually dropping off until ∼100-150 years ago, at which point the decline became much more rapid. In contrast, the central Baltic autumn spawners (CBAS) exhibited a strong population increase ∼600 YBP and continued to climb until ∼100 YBP at which point they rapidly collapsed. Coincident with the initial decline of WBAS, the central Baltic spring spawning herring (CBSS) began to increase. They plateaued around 500YBP and then started to decline ∼400YBP. Spring-spawning gulf herring (GSS) showed a strong population increase until very recently, only showing a decline in recent generations. Each population is plotted with relevant historical events noted, including the onset of particular fisheries as well as the duration of two key past climatic events: the Medieval Climate Anomaly (MCA) and the Little Ice Age (LIA), during which the Baltic was, respectively, warmer and much colder than today, albeit with short-term variability^31,56^.

**Figure 3.**
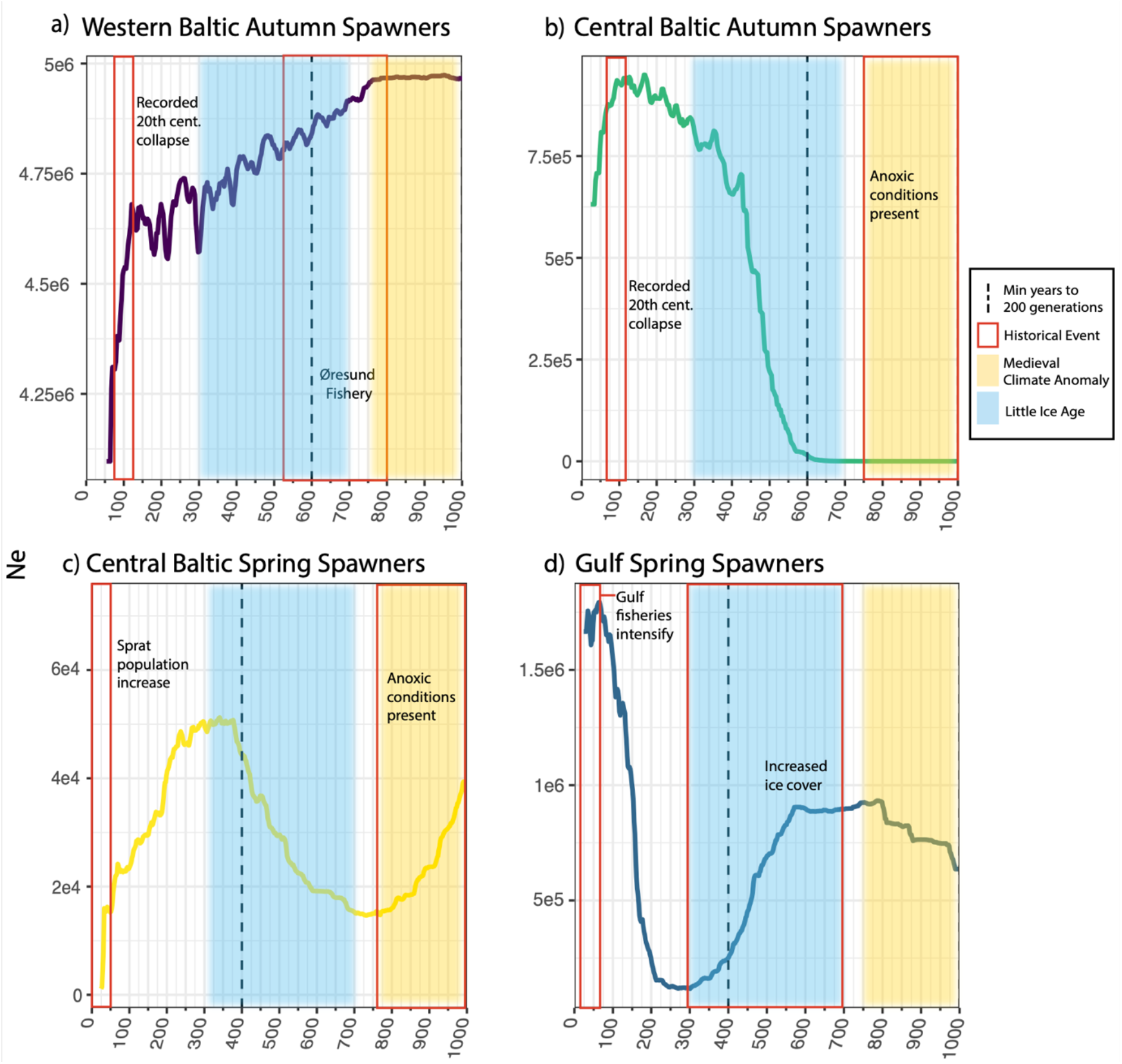
Temporal reconstruction of recent effective population sizes (N_e_) in four Baltic herring stocks. Each stock is represented in a separate chart: western Baltic autumn spawners (WBAS); central Baltic autumn spawners (CBAS); central Baltic spring spawners (CBSS); gulf spring spawners (GSS). Colored rectangles indicate key historical events. Yellow rectangles show the approximate duration of the Medieval Climate Anomaly (MCA) and blue rectangles the Little Ice Age (LIA). The dashed vertical lines show the minimum date (YBP) at which 200 generations in the past is reached (calculated using minimum generation times of 3 years for autumn spawners and 2 years for spring spawners), the known accurate window for *gone*. Historical events are denoted by red boxes for each population. a) Demographic trajectory of the western autumn spawners shows a decline starting shortly after the start of the Øresund herring. They show an additional severe decline corresponding to the reported 20^th^-century autumn spawning fishery collapse in the Baltic; b) Demographic trajectory of central autumn spawners, which appear limited during the MCA when anoxic conditions are present in the central Baltic. They increase during the LIA and rapidly decline during the period of known autumn spawning population collapse coinciding with the increase of the sprat (*Sprattus sprattus*) population; c) Central spring spawners show an increase around the time of the decline of the western autumn spawners, then a decrease again at the end of the LIA as well as another dramatic decrease around the time of the autumn spawners’ collapse ∼100YBP; d) Gulf spring spawners decrease during the LIA and then increase dramatically at the end of the LIA, starting to decline only in very recent generations when fisheries in the gulfs intensify.

## Discussion

Using modern and historical whole genome sequences from Atlantic and Baltic herring, we resolve herring population structure, reveal the earliest known large-distance herring trade and show that large-scale fishing operations had a larger impact that previously understood. We here contextualize our results with historical, ecological, and archaeological perspectives to situate our findings in a wider narrative of human-ocean interaction.

### Population assignment

All pre-20^th^ century Baltic fisheries under consideration appear to have focused on autumn-spawning herring. This observation provides genetic evidence for historical arguments that autumn-spawners were the dominant metapopulation in the Baltic in the Middle Ages. We further show that both past and present Baltic fisheries focused on Baltic herring rather than Atlantic herring as previously supposed^7^. We find no evidence of spring-spawners in the archaeological specimens analyzed. There are multiple reasons this might be the case. Historical evidence shows that the medieval fisheries, including the Øresund fishery were often strictly regulated to target spawning aggregations, and thus a single herring stock^6,26^. Further, when population sizes of the autumn-spawning herring were high, it is likely that the spring spawning herring were low; data from other oceanic basins show co-existence of the two metapopulations, but that one seasonal spawner appears to dominate^23,24^. We do not observe spring spawners in any archaeological assemblage.

Our archaeological specimens exhibited variation in adaptation to salinity. This result reflects the differentiation between herring that spawned within the Baltic proper and those that spawned within the transition zone of the Kattegat and western Baltic^36^. The higher-salinity adapted individuals appear in multiple sites in Poland, which would not have had high-salinity adapted communities spawning near the coast. Salinity conditions in the Baltic have changed slightly in the last 1000 years^57^, but not significantly enough that ocean-level salinity conditions would be present in the central Baltic. Strikingly, our samples from Truso, which date between 800-850 CE^19^, show the presence of high-salinity adapted herring with Baltic-type inversions on chromosome 12 which could only stem from fishing operations in the Kattegat or western Baltic^36^. Previous studies have shown that while there is currently a small degree of connectivity between the central and western Baltic, the majority of this is unidirectional towards the western Baltic^43^. Further, populations from the western Baltic have been shown to be genetically distinct from the central Baltic^42^, therefore it is highly unlikely that these herring were fished locally. Assemblages with later dates show a shift towards lower representation of high-salinity adapted herring (Fig 2). This reflects the decline of the Øresund fishery, and shows the existence of refugia for the Baltic autumn spawning herring metapopulation.

The discovery of high-salinity adapted herring at Truso provides genetic evidence that this assemblage stemmed from a commercial product, a hypothesis suggested by the type of bones retrieved^16^. These fish thus show the existence of long-distance fish trade in the Baltic at an earlier date than previously believed. The origin of these samples is likely in the western Baltic region, perhaps stemming from the early herring fishing operation at Rügen^10^. This observation expands the scale of earlier observations of long-distance cod trade during the Viking Age^58^ and its scope, given that herring trade is technologically more complex (requiring salting rather than only drying)^5^.

### Demographic Analysis

Demographic reconstruction showed distinct trajectories for each of the four populations under consideration. These reconstructions highlight the importance of management based on biological units, and indicate the differential effects of fishing pressure and climatic change on specific populations of the same species within the same region. We here evaluate possible varying impacts of fishing pressure and climatic change (e.g. sea surface temperature) on each of the four stocks in turn to illustrate this point.

#### Western Baltic Autumn Spawners

The WBAS exhibit a population decline coincident with the intensification of fishing in the Øresund region. High temperatures in the central Baltic have a limiting effect on autumn spawning herring reproduction^40^. We would therefore expect the end of the Medieval Climate Anomaly (MCA) (1000-800YBP) to result in increased population size for the WBAS in the absence of fishing pressure. In contrast, the WBAS begin to decline ∼800YBP and continued to decline until the present day. We observe a small drop corresponding to the proposed collapse in the 16^th^ century described by Lehmann et al.^6^, yet the population did not really contract until ∼100YBP. We conclude that the 16^th^ century decline of the Øresund fishery was therefore likely a result of local extirpation rather than a complete population collapse, yet one that clearly had a significant impact on the overall size of the WBAS stock.

#### Central Baltic Autumn Spawners

CBAS exhibit extremely low effective population size until the onset of the LIA (∼700 YBP), at which point they increase dramatically until ∼100 YBP. Sediment cores have shown that temperatures in the central Baltic during the MCA were warm enough to produce algal blooms and anoxic conditions^31,57,59^. Autumn spawning herring are more vulnerable to anoxic conditions than spring spawners as they spawn in deeper waters^60^. Combined with the temperature-limited reproductive success^40^, the CBAS were thus likely under limiting environmental conditions prior to the LIA. Our results show an expansion of this population that mirrors the timing of the WBAS decline and the onset of the Little Ice Age (LIA), a time at which there would also have been low fishing pressure in the central Baltic. Subsequently, we see a dramatic reduction in CBAS effective population size corresponding to the historically documented autumn spawning fisheries’ decline in the early 20^th^ century^25,28^.

#### Central Baltic Spring Spawners

CBSS show dramatic changes in effective population size over time. We expected to see an increase in the CBSS population concurrent with the autumn spawning decline ∼100YBP, as proposed by the hypothesis that a demographic transition occurred in the Baltic, yet this is not reflected in the results. It is unclear why the CBSS show a decline at the end of the LIA, although this is perhaps in response to the growing CBAS population. One would expect that CBSS would fill the niche left by the collapsing CBAS population in the 20^th^ century, yet it does not appear to do so. Interestingly, sprat (*Sprattus sprattus*) operate in the same niche as Baltic herring and the two species have been shown to limit each other’s population sizes^61^. Prior to the mid-20^th^ century, sprat was rare in the fishing record in the Baltic^62^, only appearing coincident with the decline of the CBAS herring. CBSS may therefore have been limited in taking over the niche left by the autumn-spawners’ decline due to an increase in the sprat population around the same time^63^. Thus, the domination of spring spawners in the 20^th^ and 21^st^ century fisheries may reflect not a population expansion of the spring spawners, but a population collapse of the WBAS and CBAS. Since the industrial fishery began targeting the CBSS ∼100YBP, we see a significant reduction in effective population size.

#### Gulf Spring Spawners

Gulf spring spawners have been subject to low fishing pressure throughout history^64^, therefore their demographic trajectory is likely more determined by changing environmental factors than the other herring stocks. Low temperatures and increased ice cover in the gulfs has been shown to have a limiting effect on gulf herring recruitment^39^, therefore a rise in population size after the LIA is to be expected and we see this in our demographic reconstructions. Fishing pressure on the gulf herring was not significant until the latter half of the 20^th^ century^64^. Our results follow this trend, in that GSS start to decline in recent generations, but it is possible that demographic changes could also be due to climate change and increasing temperatures in the gulfs. It is likely too early to tell which factors are the driving components in the changed demographic trend for gulf herring in recent generations.

#### Limitations

While *gone* is stated as accurate for a minimum sample size of 2 individuals, it is possible that small sample sizes can affect the demographic calculations^55^. Additionally, we offset each trajectory by minimum generation time for each stock due to the difficulty of accounting for changes in year-class strength over time. This is likely a conservative estimate, as herring are known to spawn in overlapping generations and both local climatic conditions^65,66^ and fishing pressure^67^ have been demonstrated to impact generation time and life history in herring. Further, *gone* is known to be highly sensitive to population structure, therefore connectivity between populations and/or the existence of substructure can affect results^55^. Given the possibility of connectivity between GSS and CBSS^43^, we further assessed the demographic trajectory of the Baltic spring spawners as a single metapopulation (Fig S10). This estimate shows a mixture of the results from the CBSS and GSS analyses, but is consistent with our conclusion that changing temperatures and fishing pressure have impacted Baltic herring demography. The combined analysis also shows a decline similar to the GSS in recent generations.

### Human Impacts on the Baltic Herring

We here present evidence for a pattern of serial depletion in the Baltic, in which the largest, most valuable herring populations were sequentially fished to exhaustion. To address growing demand for marine fish protein in Europe, the initial fishery (beginning by 850 CE and expanding especially ∼1200 CE) focused on the most economically relevant herring with the biggest spawning aggregations, the western Baltic autumn spawning herring. Once that population was exhausted, fishing pressure turned to the next-largest herring stock, the central Baltic autumn spawners and the pressure again intensified until they, too, collapsed. After the highly-valued autumn spawners were fully exhausted in the early 20^th^ century, only then did the herring fishery turn to targeting spring spawners, the smaller, less-valued population. And finally, the gulf spring spawners, long disregarded as potential commercial targets, are now facing the pressures of industrial fishing^68–70^.

Baltic herring are therefore currently facing the combined challenges of overfishing, eutrophication, rising temperatures, competition from expanding sprat populations, and further dilution of the Baltic by freshwater^9,13,23,61,71–73^. In recent years, ICES have recommended a MSY of 0 for multiple Baltic herring stocks^15,15,68,74,75^. Yet, fishing continues and the stocks do not recover^76^ because economic and cultural factors continue to drive exploitation. In the face of these issues, it may seem that the fate of the herring is sealed. However, modeling of Baltic ecosystem dynamics has shown that fisheries policy and conservation management are some of the most important factors in determining the long-term sustainability of the Baltic^977^. Therefore, it is crucial that we learn from the past, both recent and further-reaching, to better understand the real impact of various management and extraction policies on the demography of Baltic herring throughout time. By bringing historical ecology methods into play, we here elucidate ecological and/or cultural thresholds^78^ that were crossed as the Baltic fisheries changed over time, finally resulting in the Baltic we see today. These thresholds – including cultural change (increased demand for marine fish), exploitation levels, and ecosystem change – can be used to inform better management policies in the future by reframing what constitutes healthy population dynamics (size, migratory behavior, etc.) to better reflect long-term ecosystem dynamics.

The gulf spring spawning herring are the last remaining healthy herring stocks in the Baltic^79,69^. They are also the stock most adapted to the brackish conditions which are expected to increase in the Baltic into the future. In order to preserve this stock, it is crucial that management bodies understand the pattern of serial depletion in the Baltic, thereby providing avenues to identify specific exploitation strategies that have resulted in population collapse in the past^80^ and avoid these for the future. The demographic trajectories of the central Baltic autumn spawners and the gulf spring spawners provide further avenues of hope for the Baltic herring. Both stocks’ demographic reconstructions indicate that in the absence of industrial fishing pressure, herring stocks can rapidly recover from long-term exposure to poor environmental conditions. If the changing climate in the Baltic can be addressed well enough to maintain suitable threshold conditions for herring and fishing reduced in line with ICES recommendations, we may yet be able preserve a significant spring spawning Baltic herring population.

## Materials and Methods

### Archaeological Material and Laboratory Methods

Archaeological bone samples were obtained from 7 sites in Poland, Denmark, and Estonia spanning 800-1600 CE (Fig 1A). A full sampling table with site information can be found in Supplementary Dataset S1. Samples were processed following the laboratory pipeline in Ferrari & Atmore et al.^50^. Full laboratory methods can be found in Supplementary Information, with library protocols detailed in Supplementary Dataset S1. All laboratory protocols were carried out in the dedicated ancient DNA laboratory at the University of Oslo following regular decontamination and authentication protocols^81–83^. Samples yielded 0.002-2.5X coverage.

### Modern Genome Sampling

53 tissue samples were collected around the Baltic and the Norwegian coast between 2002 and 2010 as reported in Ruzzante et al.^84^ and André et al.^85^ (Fig 1B). DNA was extracted from the tissue samples using a DNEasy Blood and Tissue Kit^86^. Library build and sequencing was carried out at the Norwegian Sequence Centre. Samples were sequenced on an Illumina HiSeq 4000, yielding coverage of 7-17X. Full metadata for these samples can be found in Supplementary Dataset S2. An additional 22 sequences were obtained from previously published data to ensure that all major herring populations were represented^36^. Coverage from publicly available sequences ranged from ∼15-55X.

### Alignment and SNP Calling

Both modern and ancient sequences were aligned to the herring reference genome Ch_v2.0.2^35^. All sequences were aligned following the PALEOMIX pipeline^87^. Modern sequences were aligned using bwa-mem and ancient sequences were aligned using bwa-aln. mapDamage2.0^88^ plots for post-mortem deamination were assessed to validate the ancient samples (Fig S1). Modern nuclear sequences were further processed following the GATK best practices pipeline with GATK4^89^. Full methods are described in the Supplementary Information. Several individuals from the modern dataset were removed due to suspected contamination and/or incorrect metadata from the publicly available dataset. The final modern dataset comprised 52 newly-sequenced genomes and 16 publicly-accessed genomes. The process of data cleaning and individual assessment is detailed in the Supplementary Information (Figs S2-S5).

### Determining Population Structure

smartPCA^90,91^ (Eigensoft v7.2.1) was run on the entire modern nuclear sequence dataset to assess population structure. A maximum-likelihood phylogenetic tree was then built with IQTree^51,52^ using the mitogenome dataset including all modern and ancient samples to verify that the archaeological samples are Atlantic or Baltic herring. A Pacific herring (*Clupea pallasii*) mitogenome – obtained from Han et al.^36^ – was used as an outgroup. Genetic diversity (*π*) was estimated in 100kb windows along the nuclear genome using VCFtools for each of the 4 Baltic populations: western Baltic autumn spawners (WBAS), central Baltic autumn spawners (CBAS), central Baltic spring spawners (CBSS), and gulf Baltic spring spawners (GSS) with 2-3 samples representing each population. Full metadata for modern sequences can be found in Supplementary Dataset S2. Fine-scaled population structure was determined using KING^92^, which calculates individual pairwise kinship coefficients within each population. Baltic spring spawners and Baltic autumn spawners were assessed as metapopulations to determine the level of relatedness between possible subgroups (e.g. gulf vs central Baltic spring spawners and western vs central Baltic autumn spawners).

### Population assignment

*BAMscorer* is capable of accurate genomic assignments using extremely low-coverage data^50^. Previous studies have reported that population structure in herring is driven by spawning season and adaptation to different levels of salinity^22,33–36^. Three sets of assignment tests were designed for *BAMscorer* for the following assignments: spawning season, chromosome 12 inversion type (Atlantic vs Baltic), and adaptation to salinity. The spawning season database was built using previously-published diagnostic loci, as was the salinity adaptation database^36^. Following the methods from Ferrari & Atmore et al.^50^ for database creation and parameter testing, a database with 4503 SNPs in the chromosome 12 inversion was created using the default parameters from the *BAMscorer wrap_snps* module. Full methods and results of assignment sensitivity analysis can be found in the Supplementary Information. Each test was then applied to ancient alignment files with sufficient reads for accurate assignment.

### Demographic reconstruction

Runs of homozygosity (ROH) were analyzed per population to determine the general trends in effective population size for each population (complete methods can be found in the Supplementary Information). The timing of the coalescent events associated with specific windows of ROH length were determined using the formula *L = 100/2g cM*^93^ where *L* refers to the length and *g* is generation time (Fig S11). ROHs were subsequently binned into 2 groups corresponding 650-400 YBP and 400-200 YBP. We compared the total sum and number of ROH for each bin as well as across all bins.

The four Baltic datasets were input into the demographic software package, *gone* ^55^, which estimates Ne in the recent past using LD decay. Ne estimates are geometric means taken from 40 bootstrapping iterations that randomly sample 50,000 SNPs from each chromosome in each population to estimate LD decay. *gone* has been shown to be robust to natural selection^94^ and viable for small sample sizes^55^. No MAF filter was applied to the dataset used for *gone*. Default parameters were used except for substituting the known population-wide recombination rate for Baltic herring of 2.54 cM/Mb^35^. *Gone* results were scaled by generation using the minimum generation time in herring (3 years for autumn spawners and 2 years for spring spawners^95^). Each population trajectory was additionally offset by the year the sampling took place (1979-2016, see Supplementary Dataset S2). As previous studies have suggested some connectivity between GSS and CBSS^43^, an additional analysis was undertaken using the Baltic spring spawners as a single population (Fig S16). Full *gone* methods are described in the Supplementary Information.

## Acknowledgments

We thank Agata Gondek for processing of the first herring samples as well as Giada Ferrari for guidance with lab work. We are further grateful to Pernille Bangsgaard and Kurt H. Kjær at the Natural History Museum of Denmark for their assistance with archaeological sampling.

