## Supplementary Information for "Serial depletion of Baltic herring since the Viking Age revealed by ancient DNA and population genomics"

**This PDF file includes:**

Supplementary text

Figures S1 to S16

Table S1

SI References

**Other supplementary materials for this manuscript include the following:**

Datasets S1 to S2

Supplementary Information Text

**Full ancient DNA laboratory methods**

Single-bone samples weighing 10-70mg were bathed in UV light to clean the exterior, then placed in digestion buffer (1 ml 0.5 M EDTA, 0.5 mg/ml proteinase K, 0.5% N-Lauryl sarcosine) and crushed in 1.5ml Eppendorf tubes using single-use plastic micro-pestles. DNA was extracted following the double-digest procedure from Damgaard et al.^1^ with MinElute PB buffer (QIAGEN). Samples were then purified through MinElute columns using the QIAvac 24 Plus vacuum manifold system (QIAGEN) for a final elution volume of 65ul. Two types of libraries were built, with some samples built using the double-stranded library protocol from Meyer & Kircher^2^ and some built with single-stranded libraries following the Santa Cruz protocol^3^. A full list of the library protocols used per sample can be found in Supplement S1. All laboratory protocols were carried out in the dedicated ancient DNA laboratory at the University of Oslo following regular decontamination and authentication protocols^4–6^. Each library underwent 12-15 cycles of PCR amplification followed by purification using the Agencourt AMPure XP PCR purification kit using a 1:1 bead:sample ratio. Libraries were then assessed for quality on a Fragment Analyzer^TM^ (Advanced Analytical) using the DNF-474 High Sensitivity Fragment Analysis Kit to determine suitability for sequencing. Samples of high enough quality were then sequenced on an Illumina HiSeq 4000 and/or NovaSeq 6000 at the Norwegian Sequencing Centre.

**SNP calling and filtering**

Modern samples were filtered using bcftools v1.3^87^ (*FS<60.0 && SOR<4 && MQ>30.0 && QD > 2.0 && INFO/DP<415140' --SnpGap 10*) and VCFtools v0.1.16^49^ (*--minGQ 15 --minDP 3 --remove-indels --maf 0.01*). Non-biallelic loci were removed. An additional dataset with no MAF frequency filtering was created for *gone* analyses known to be sensitive to removing minor alleles. Mitogenomes were called and filtered as above for ancient and modern samples. Individuals with >30% missingness were removed.

**Removing outliers from the whole-genome dataset of modern herring specimens**

Our dataset contains whole genome data of herring specimens obtained from a range of difference sources, collected over a number of years. The dataset further combines publicly available data with data generated *de novo* for our study. Given this wide range of sources, we performed a number of exploratory analyses to ensure data integrity. First, we previously identified that two of the individuals from the Han et al.^7^ dataset were technical duplicates^8^, supported by using KING^9^ (Table S1). One of these individuals was chosen at random to be included in the dataset and the other was discarded. Second, exploratory population analyses using smartPCA^10,11^ indicated that several individuals were significant outliers (Figure S2). To further explore this pattern, levels of heterozygosity and inbreeding coefficients were calculated on an individual basis for the modern nuclear data using VCFtools. Three individuals (HER_Z12_IsleOfMan, HER_NSSH34, and M-HER004) showed unusually high levels of heterozygosity and inbreeding coefficients (Figures S3, S4). We further assessed the possibility of contamination by analyzing levels of heterozygosity along the mitogenome (Figure S5). Again, three individuals (HER_NSSH33, HER_NSSH34, and HER_Z12_IsleOfMan) showed clear signs of contamination. Based on these results, four individuals (HER_Z12_IsleOfMan, HER_NSSH33, HER_NSSH34, and M-HER004) were removed from the dataset. Finally, one individual from the Han et al (Ref) dataset was removed as it consistently clustered with individuals with non-matching metadata (Figure S6). Given the earlier inconsistencies of the metadata, this sample was also removed. The cleaned dataset was used for demographic (runs of homozygosity, $\pi$, KING, and *gone*) and for whole-genome PCA analyses.

**BAMscorer sensitivity analysis**

In order to determine which ancient sequences could reliably be scored, we first assessed the required read depth to accurately assign each test. Required read depth per assignment was assessed following the downsampling and bootstrap method of Ferrari & Atmore et al.^8^. We selected 8 samples from the modern dataset to independently test the power of assignment probability with known metadata. The alignment files of these eight samples were randomly downsampled to between 1k and 100k reads 20 times and then used to assess BAMscorer sensitivity for each assignment test. Given the relatively low number of autumn spawning herring, we investigated whether including the removed individuals (see above) impacted on the biological patterns for each comparison. Although these outliers impact the whole genome population analyses, SmartPCA analysis on each of the three BAMscorer assignment databases showed that including these outliers associated with contamination bias did nothing to change the shape of the distributions. Due to limited sample size for the different categories in these assignment tests, several outliers were left in the BAMscorer databases apart from AAL1_CelticSea (suspected incorrect metadata) and Gavle54 (identical to Gavle98) from the Han et al.^7^ dataset.

The modern database showed strong differentiation between spring and autumn/winter spawning seasons, following previously reported results^7,12,13^. Sensitivity analysis showed that spawning season can be accurately determined in alignment files with as few as 50 000 reads (Fig S7). The chromosome 12 inversion could be confidently assigned with as few as 5 000 reads (Fig S8) using default parameters. Given that only samples for which spawning season could be assigned were analysed, all samples still retained at least 50 000 reads. Salinity scores could be determined for samples with at least 60 000 reads (Fig S9).

**Evaluating demographic independence**

Demographic independence of the Baltic subpopulations was further assessed by calculating individual pairwise relatedness for each group with KING. To assess the substructure in the Baltic, metapopulations were grouped, resulting in two Baltic populations: autumn spawners and spring spawners (for full KING results, see Supplementary Dataset S2). In the Transition Zone (TZ) population, M-HER066 showed strongly negative kinship coefficients with the rest of the TZ individuals. This is a sign that there is population structure, thus M-HER066 is likely not actually a TZ individual, but could be part of the Atlantic spring spawners population, which is where it clusters on the PCA. M-HER066 was removed from analysis to eliminate outlier bias. Kinship coefficients were plotted per population to visualize the distribution of relatedness in each population (See Fig S10).

A one-way ANOVA showed that metapopulation ID was significantly associated with kinship coefficient mean and variance (p=8.16e-14, DF=4). Baltic spring spawners had the lowest average kinship coefficient (0.0336). Baltic autumn spawners had the second highest kinship coefficient (0.05827), likely due to the effect of the Fehmarn individuals, which showed high relatedness to each other. The Fehmarn within-group average was 0.08. This illustrates there is some substructure in the herring metapopulations, although it is likely that there is a degree of connectivity, as strong substructure would result in negative kinship coefficients. It should be noted that all individuals showed low levels of relatedness and there was some variance in relatedness between all groups. The Transition Zone showed the highest variation in kinship coefficient, likely due to the presence of subpopulations such as the Idefjørd herring.

**Runs of Homozygosity**

Initial ROH results showed that the variation in ROH length, count, and total sum were largely determined by differences in sample size. Therefore, a subset of each population was chosen randomly, with 2-3 individuals per population (samples used can be found in Supplementary Dataset S1). PLINK files were generated from the modern nuclear sequence data and then assessed for differences in runs of homozygosity (ROH) using PLINK 1.9^14^ following previously-published recommendations^15–17^. The following command was used: *plink –bfile herring --chr-set 26 --double-id --homozyg-snp 50 –homozyg-kb 90 –homozyg-density 50 –homozyg-gap 10000 –homozyg-window-snp 50 –homozyg-window-het 3 –homozyg-window-missing 10 –homozyg-window-threshold 0.05 --out herring_roh.* ROHs were compared by length, count, and total sum between the 4 Baltic populations.

***gone* analysis**

We reconstructed population demography using *gone,* which is reported to be accurate up to 200 generations in the past^18^. We show trajectories up to 1000 YBP are shown here to reflect the possibility of overlapping and/or longer generation times. As herring spawn in overlapping generations with varying maturation times, and can live up to 18 years^19^, generation times in five and ten-year increments are further shown in Fig S14 to illustrate the possible variations in trajectory for each population. Exact N_e_ estimates of *gone* are likely affected by large effective population sizes in the past and the small sample size used here^18^. True N_e_ estimation is further confounded by fish spawning behaviors, including overlapping generations and batch spawning^20^. Therefore, the key findings from the *gone* analysis are the divergent trajectories and their relative changes rather than the absolute values of N_e_ estimated for each population at a given date. All four populations showed unrealistic bottlenecks to near-zero N_e_ in the four most recent generations. The original *gone* paper reports similar estimates for recent generations when using small sample sizes (n **≤** 10) and applications to other fish populations show a similar pattern^18,20^, therefore these recent extreme bottlenecks were disregarded as an artifact of the calculations and removed from the results. Given the possibility of connectivity between the GSS and CBSS populations, *gone* was also used to estimate demographic history for the Baltic spring spawning metapopulation as a whole (Fig S16).

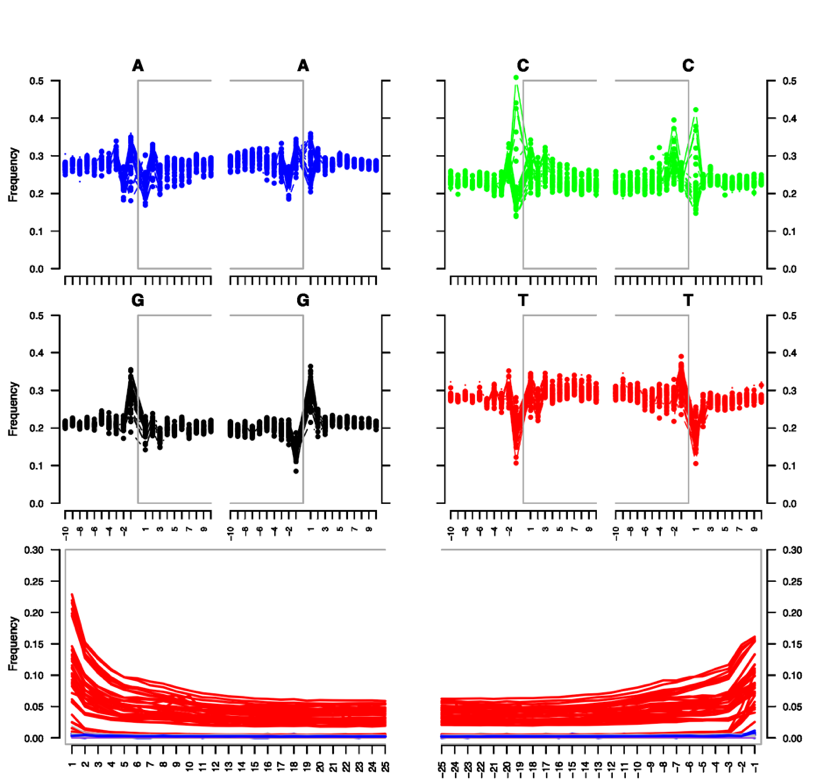

a

b

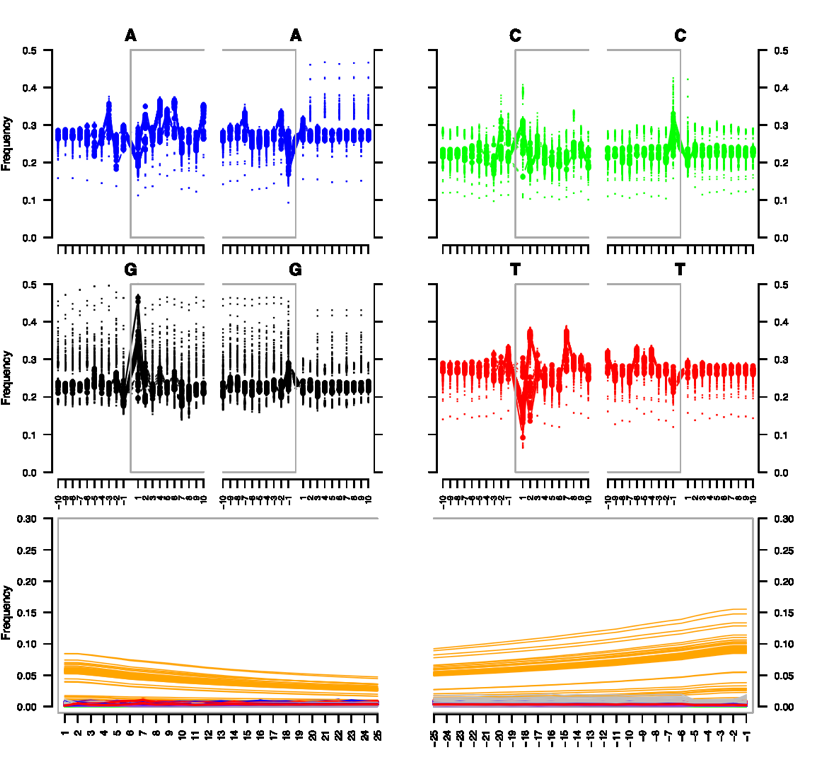

**Figure S1 – Fragment Misincorporation and Damage plots.** a) These plots show that the post-mortem deamination patterns of all archaeological herring specimens (n=40) are characteristic of those associated with ancient DNA^21^. b) Fragment misincorporation and damage plots for modern herring sequence data (n=72).

**Table S1 – KING output for duplicates/relatedness in modern herring reference data**.

Concordance values above 0.8 are indicative of duplicates according to the KING documentation. Table from Ferrari et al^8^.

| FID1 | ID1 | FID2 | ID2 | N | N_IBS0 | N_IBS1 | N_IBS2 | Concord | HomConc | HetConc |
| --- | --- | --- | --- | --- | --- | --- | --- | --- | --- | --- |
| Gavle54 | Gavle54 | Gavle98 | Gavle98 | 11314407 | 187 | 4995 | 11309225 | 0.99954 | 0.99998 | 0.99674 |

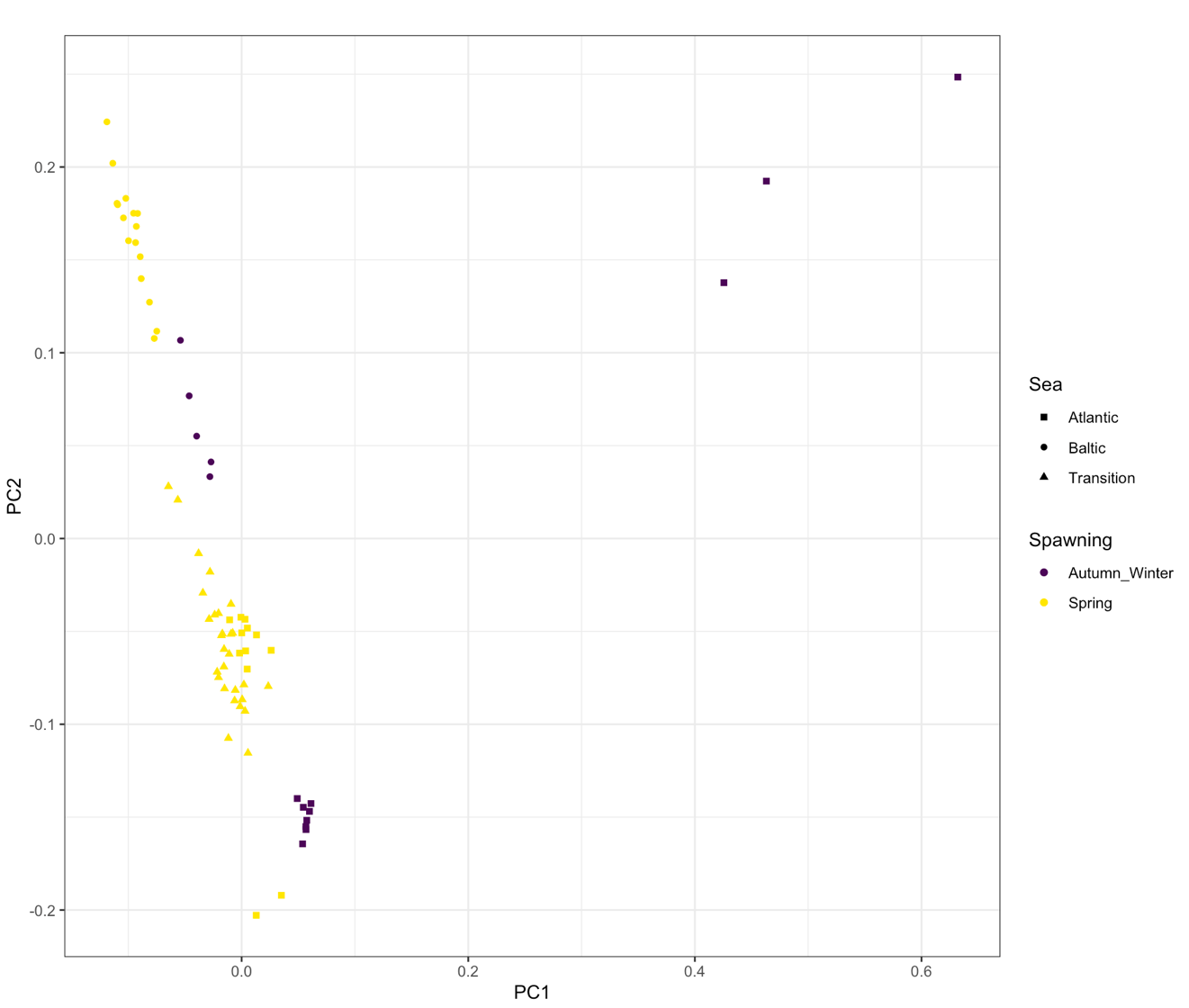

**Figure S2 – Exploratory population analyses of Atlantic herring using genome-wide data.** The PCA is based on 10,368,446 SNPs. The color indicates the spawning season and the shape indicates the sea in which the sample was collected. “Transition” refers to the area between Norway, Sweden, and Denmark that spans the transition between the North Sea and the Baltic Sea. Three Atlantic herring specimens are located away from the main herring clusters. This pattern is driven by inclusion of a single specimen (HER_Z_12_IsleOfMan) that is contaminated (see also Figure S3, S4, and S5).

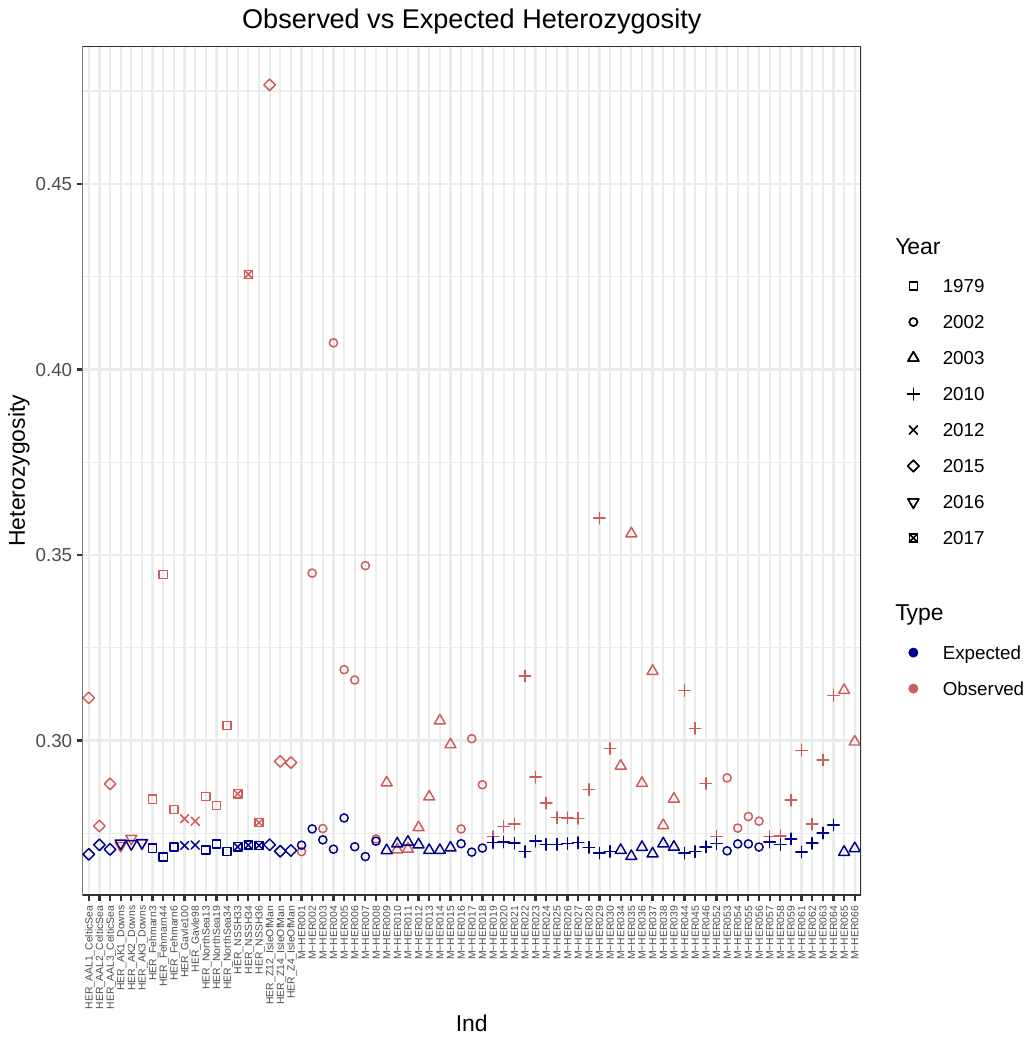

**Figure S3 – Observed vs Expected heterozygosity in modern herring specimens.** Individual herring specimens are ordered along the x-axis. Expected heterozygosity (blue) and observed heterozygosity (red) vary from individual to individual. Shapes indicate the year in which the sample was obtained. Three individuals are clear outliers – HER_Z12_IsleOfMan, HER_NSSH34, and M-HER004.

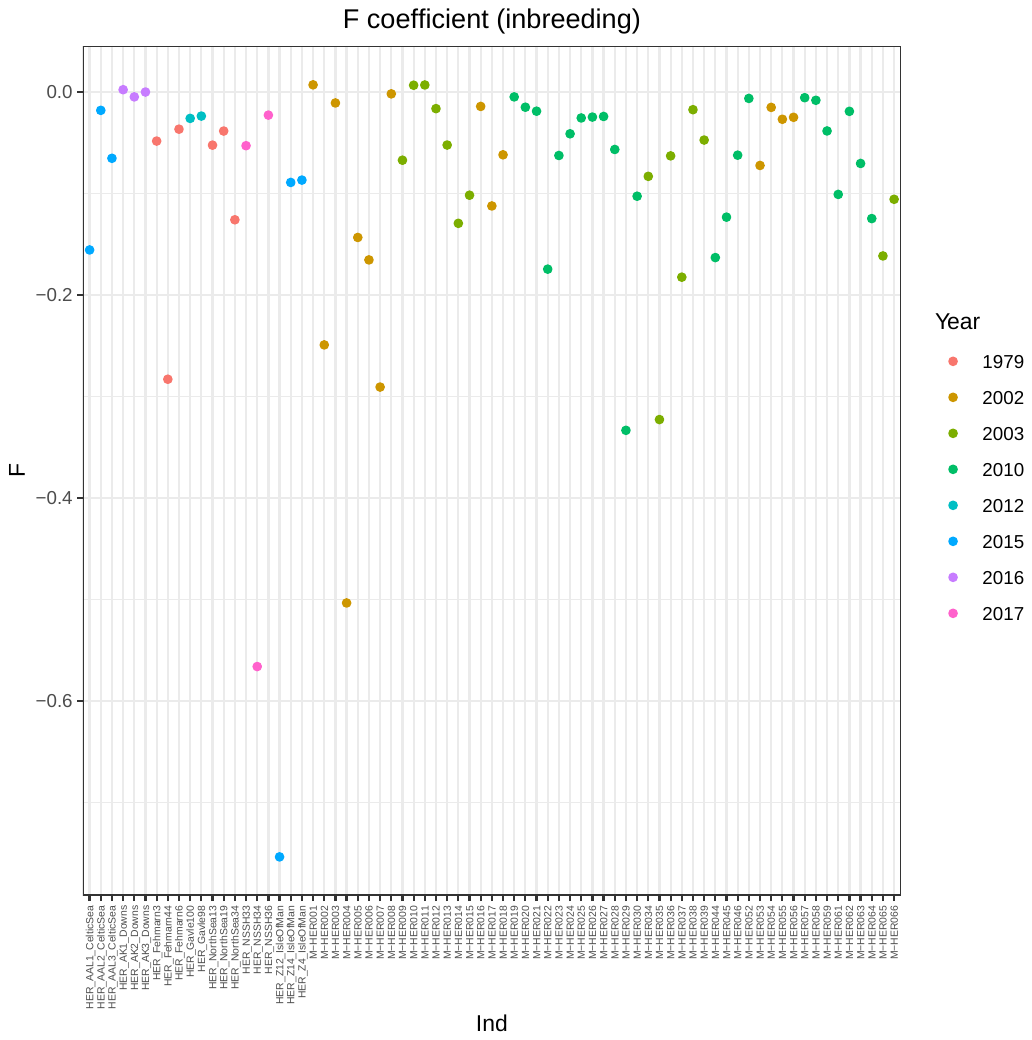

**Figure S4 – F coefficient in modern herring sequences.** Individual herring specimens are ordered along the x-axis. Plot of F coefficient color-coded by year the sample was obtained. Three individuals have extremely low F coefficients. These are the same individuals (HER_Z12_IsleOfMan, HER_NSSH34, and M-HER004) as in Figure S3.

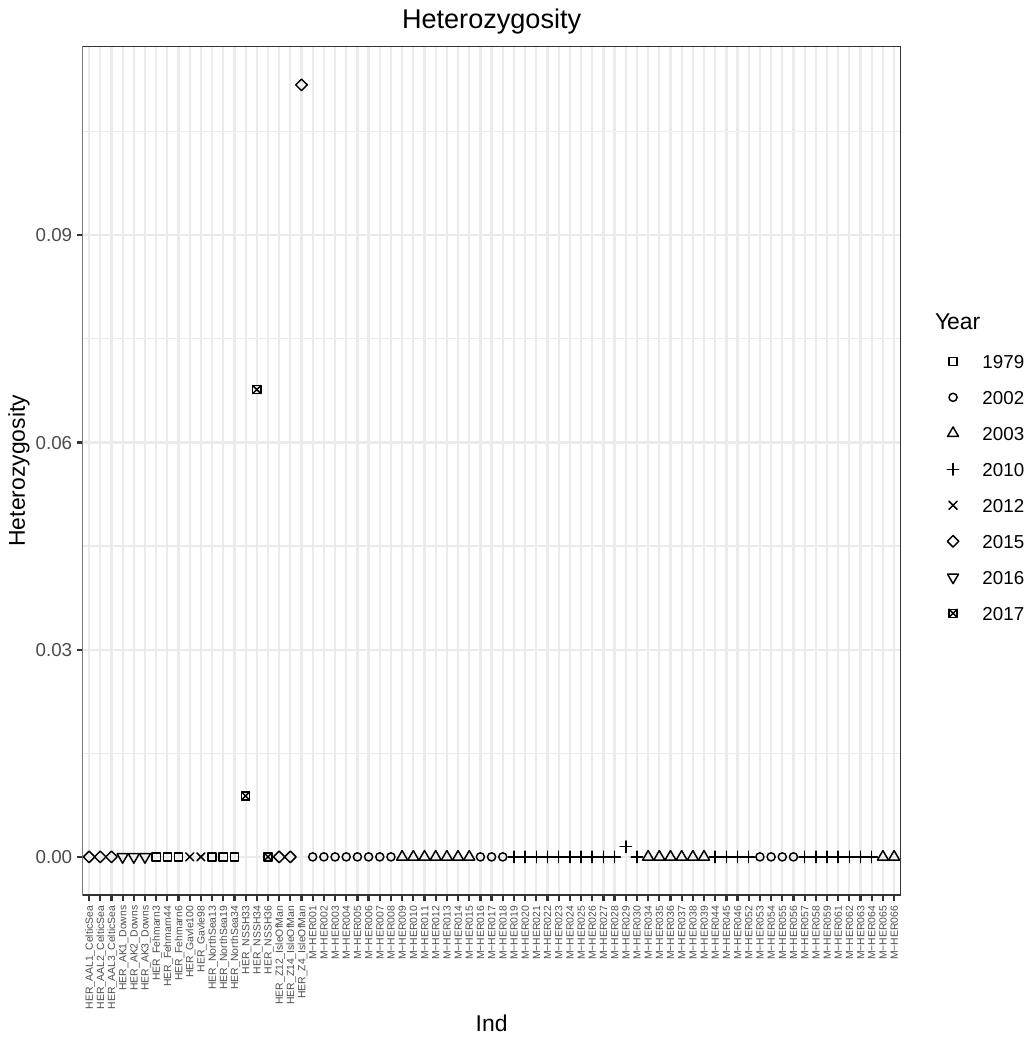

**Figure S5 – Levels of heterozygosity observed in the mitogenome of modern herring specimens**. The mitogenome was coded as diploid and heterozygosity was calculated using VCFtools. The three individuals with heterozygosity values >0 are indicative of contamination. These are HER_NSSH33, HER_NSSH34, and HER_Z12_IsleOfMan. M-HER004 does not exhibit signs of contamination in the mitogenome, but was still removed from the dataset due to its high levels of heterozygosity and low values of the F coefficient.

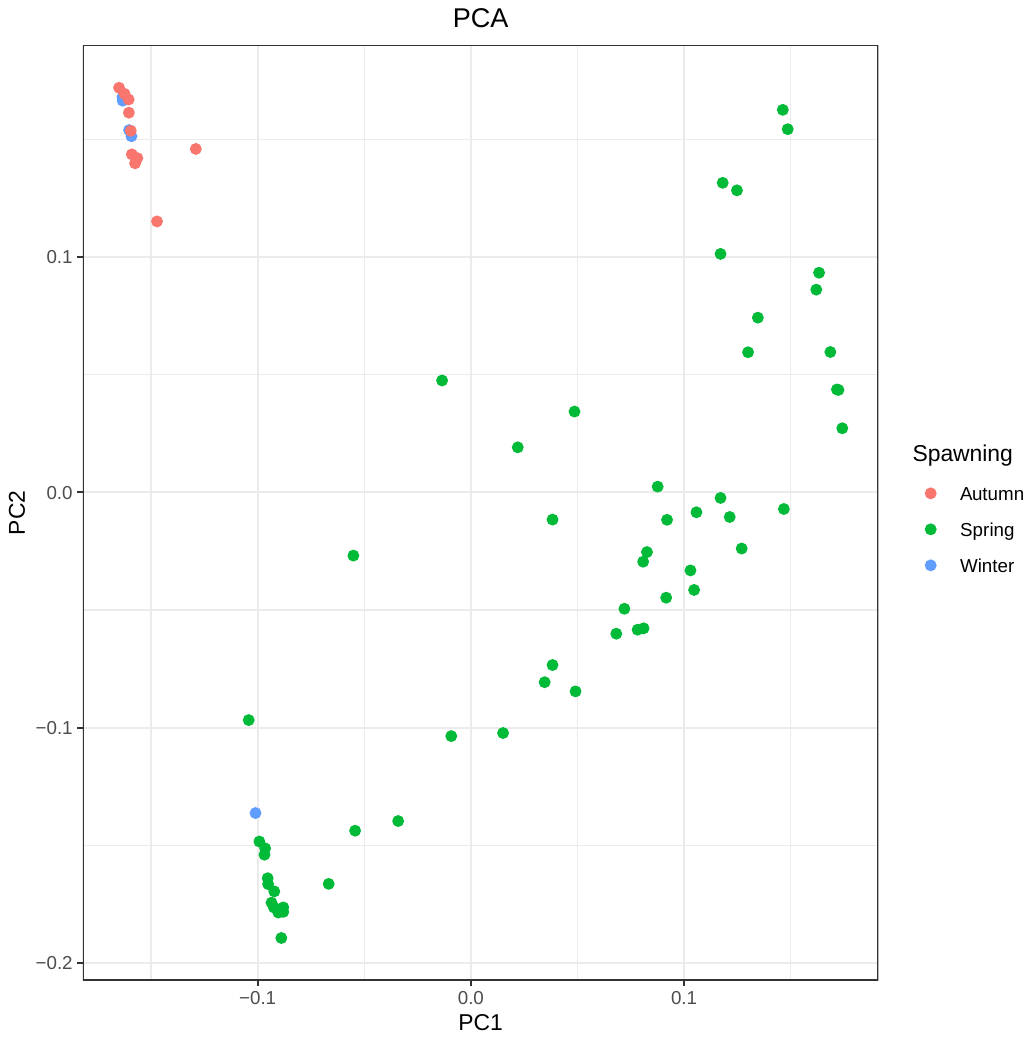

**Figure S6 – PCA of spawning loci including all modern herring specimens.** Spawning loci show a clear distinction between spring spawners and autumn/winter spawners. HER_AAL1_CelticSea from the Celtic Sea is coded as a winter spawner but consistently clustered with spring spawners in smartPCA analysis.

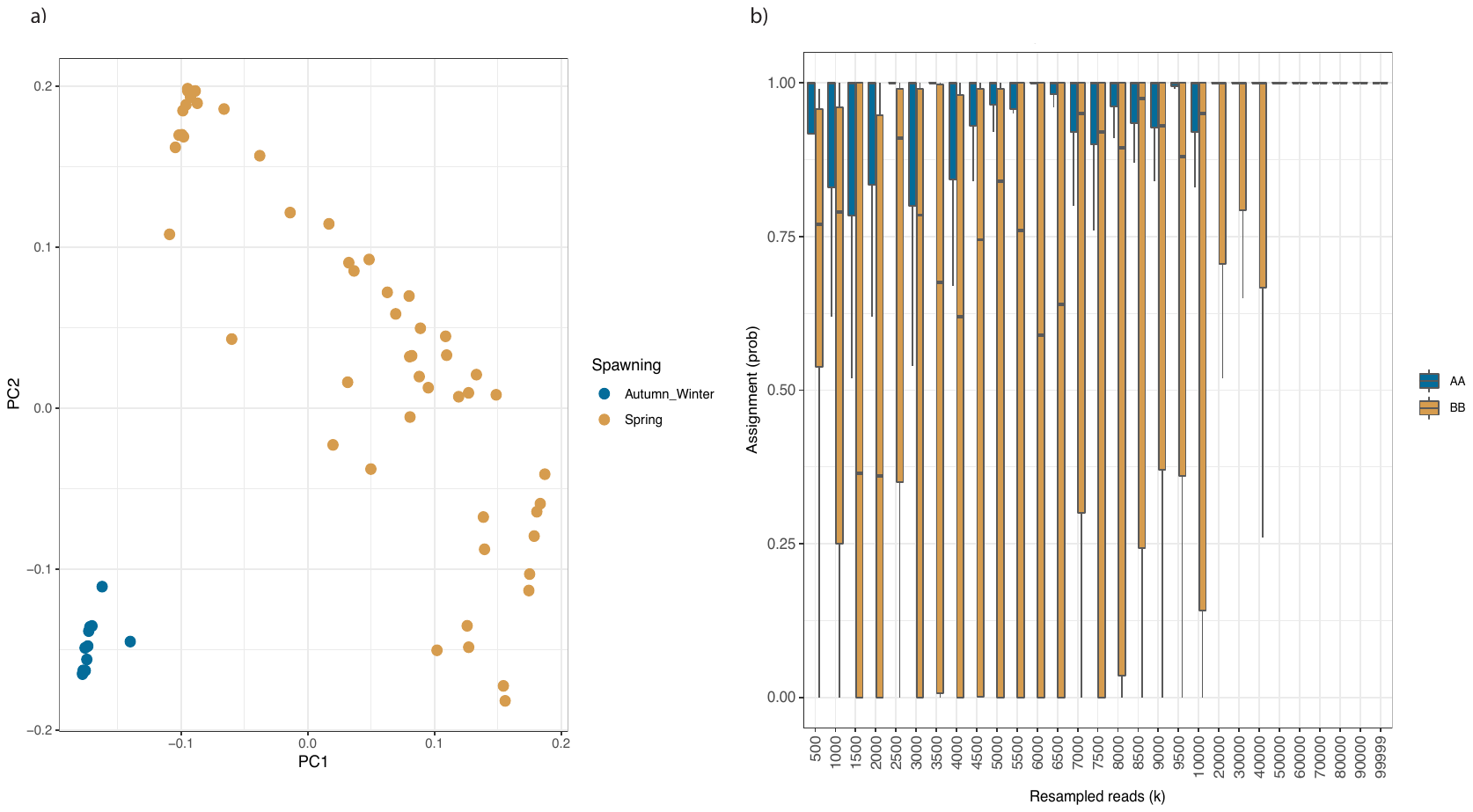

**Figure S7 – Spawning season assignment sensitivity.** a) PCA of the diagnostic loci for identifying spawning season from Han et al.^7^; b) Application of BAMscorer to the bootstrapped downsampled alignment files with known metadata show greater difficulty assigning spring spawning than autumn spawning. Both seasons can be assigned with no error at 50 000 reads. AA type is autumn spawning and BB type is spring spawning.

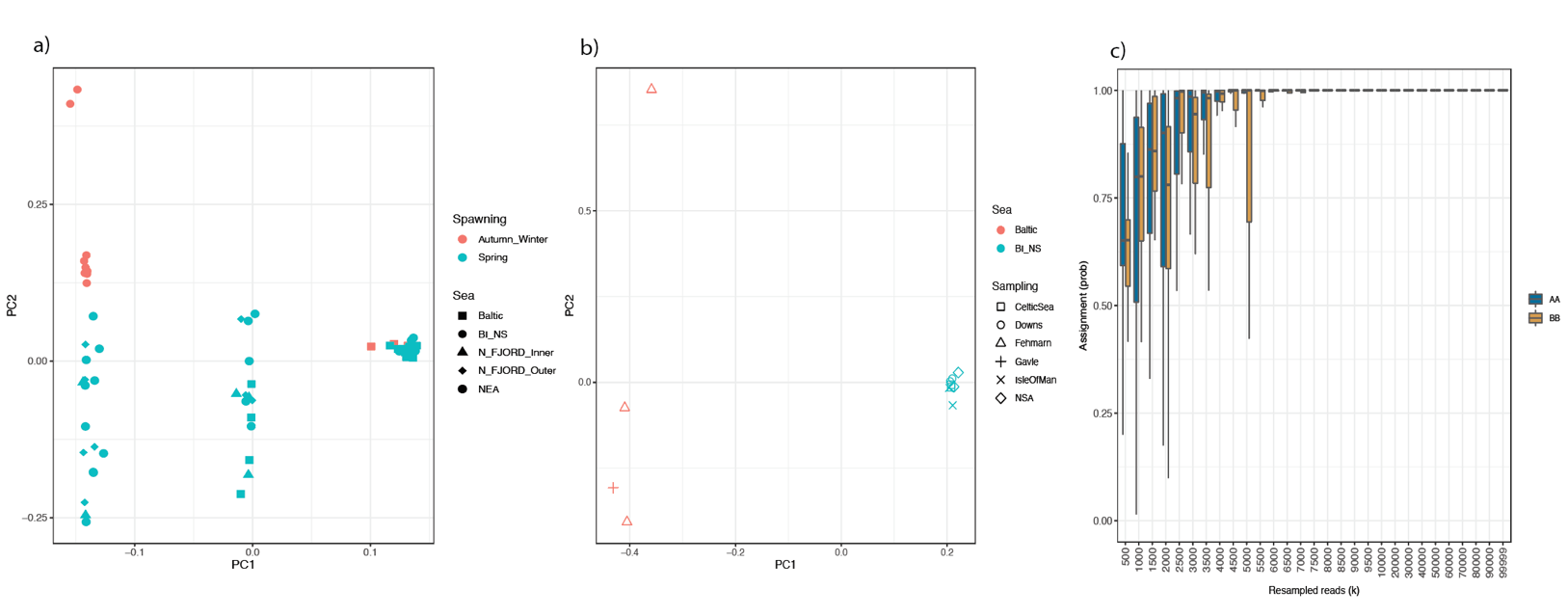

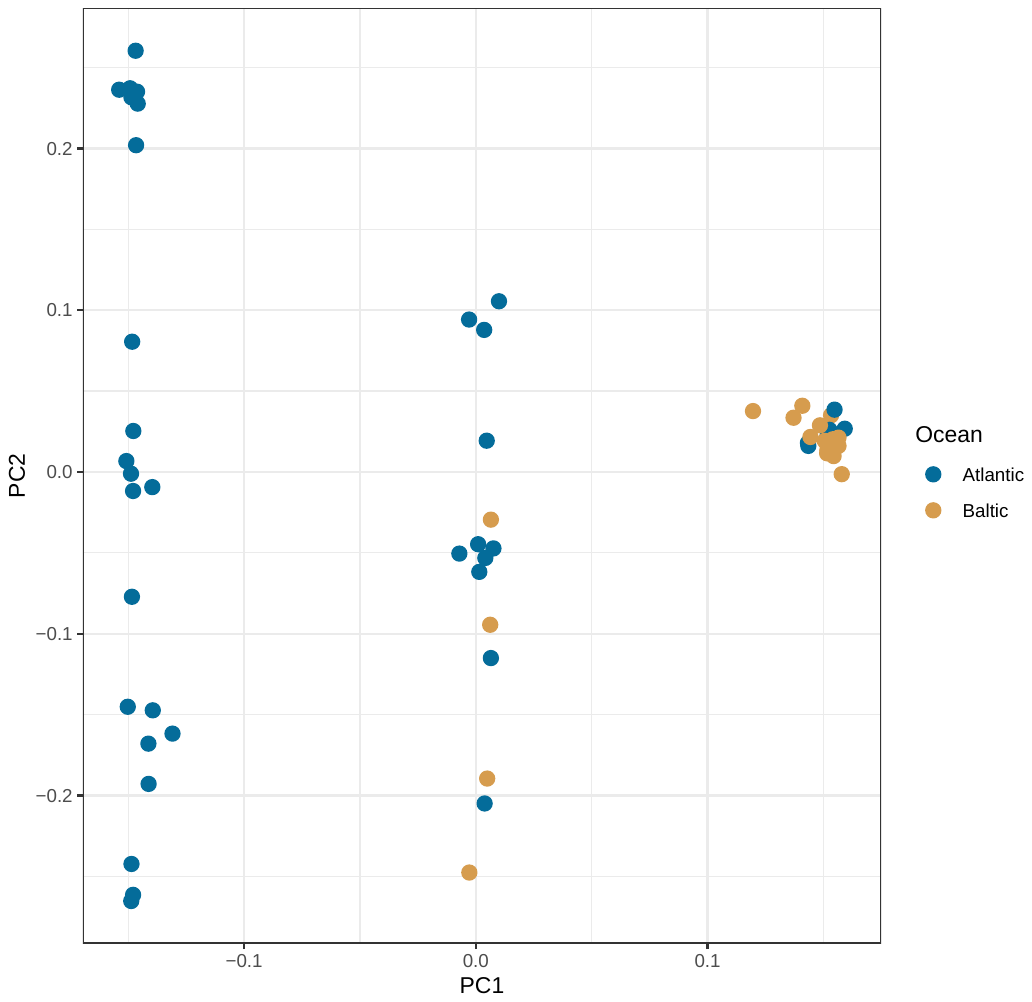

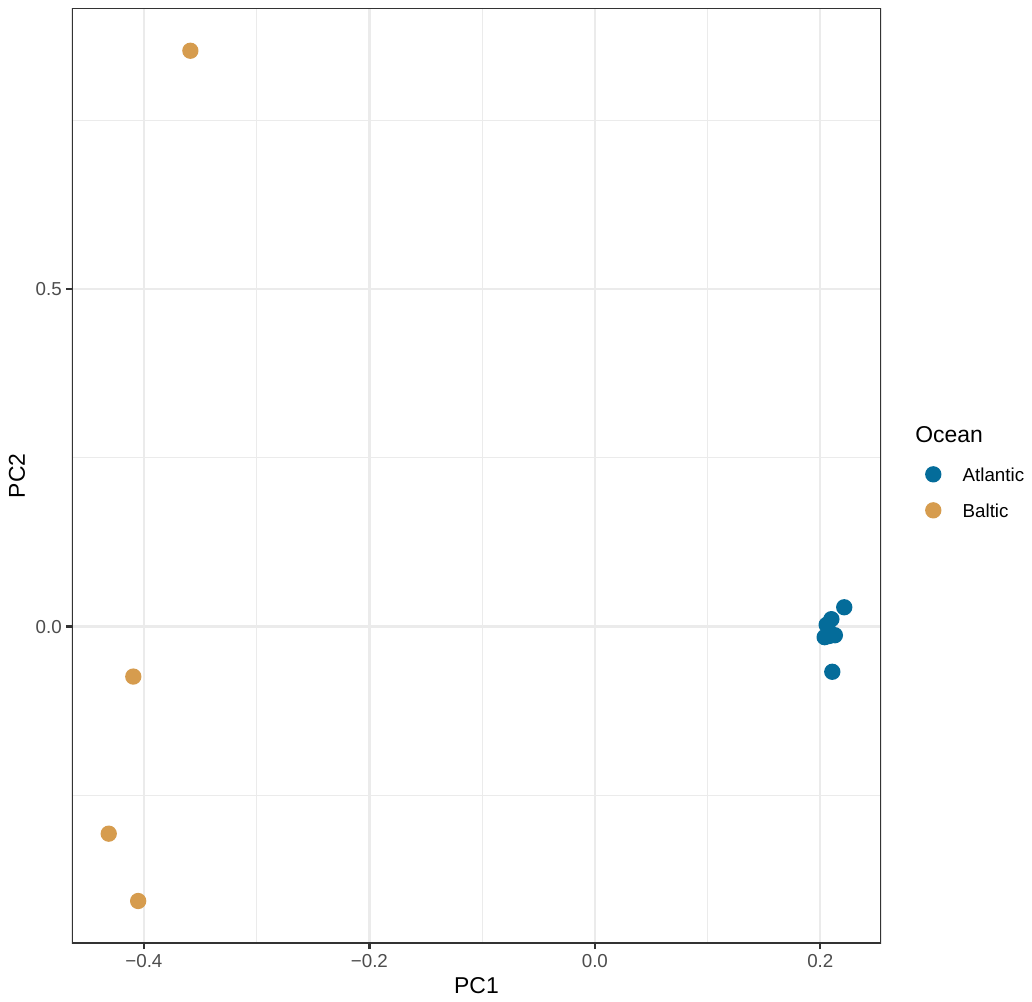

b)

a)

**Figure S8 – Chromosome 12 inversion split between Atlantic and Baltic haplotypes.** a) Chr12 inversion for all individuals in the modern dataset. The spring spawners exhibit no structure at this locus associated with geography, which confirms the results reported by Han et al.^7^. b) The autumn spawners show a split between the Atlantic and Baltic type inversions. c) The inversion locus can be scored with as few as 5000 reads.

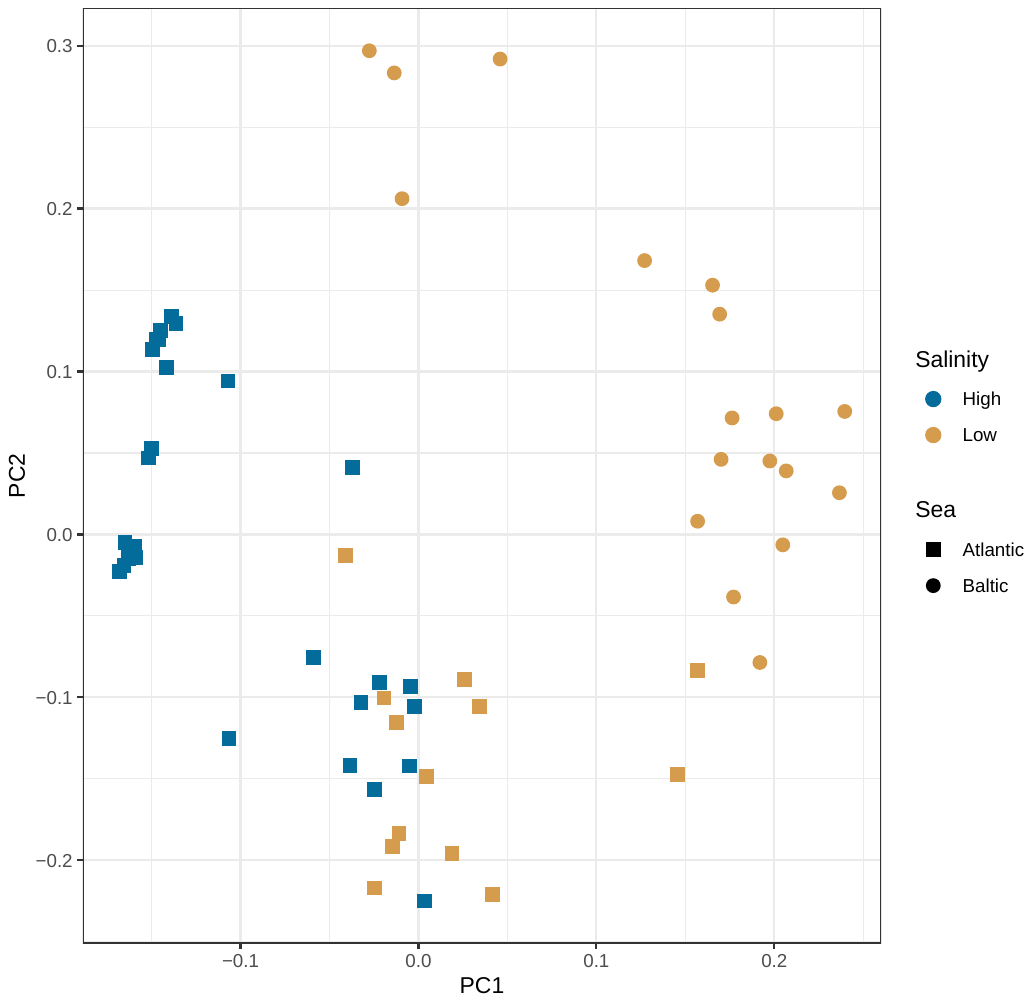

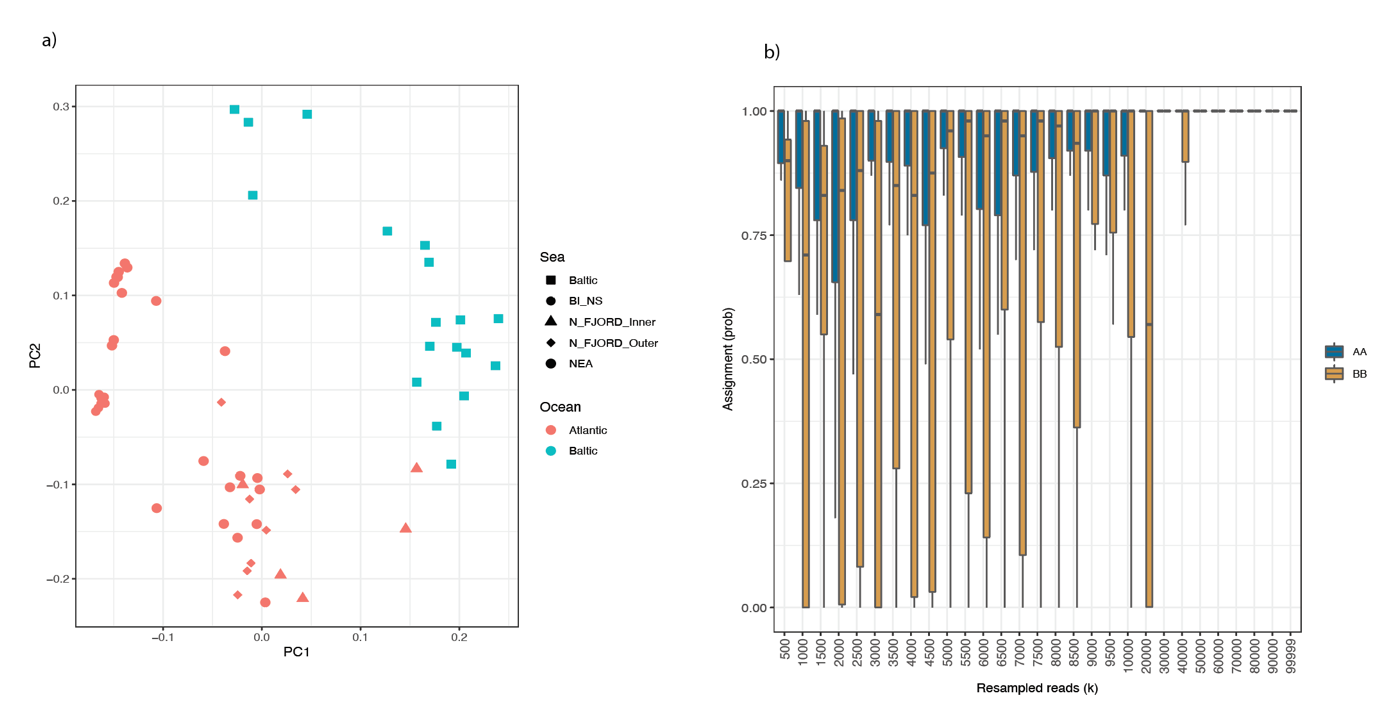

a)

**Figure S9 – Salinity adaptation loci BAMscorer database creation and parameter testing.** a) PCA generated with smartPCA. Color indicates ocean basin as a proxy for salinity (high being Atlantic, low being Baltic) and shape indicates population of origin. Some Atlantic samples were from Norwegian fjords with low salinity therefore they fell between the two groups. These samples were removed from the dataset for the final scoring. b) Iterative application of BAMscorer to downsampled alignment files showing a minimum of 50,000 reads is required for determining salinity adaptation.

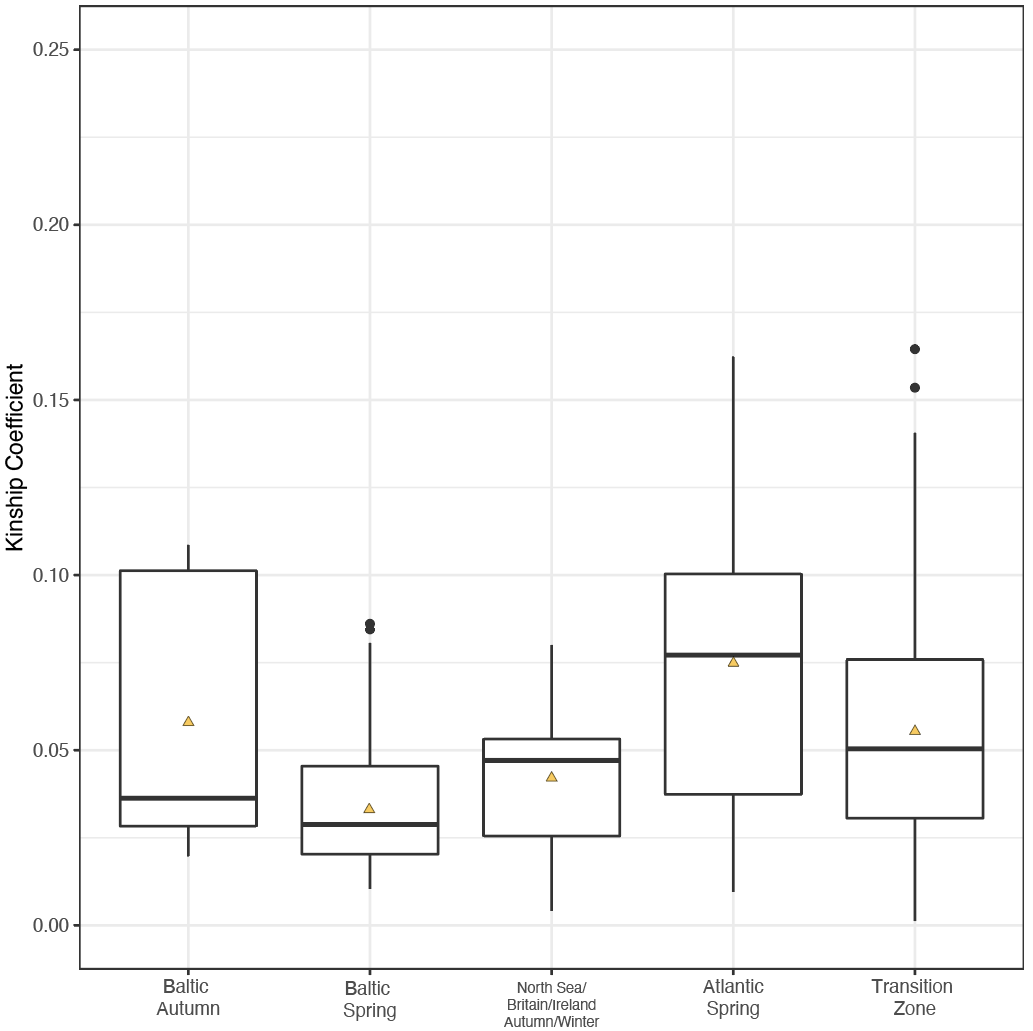

**Fig S10 – Individual pairwise relatedness by metapopulation.** Each boxplot shows the distribution of kinship coefficients across the metapopulation along the x-axis. Baltic autumn and Baltic spring populations were grouped as two metapopulations to assess the level of substructure in each population; a lower kinship coefficient is indicative of higher substructure. Lines indicate median kinship coefficient and yellow triangles indicate the mean kinship coefficient per metapopulation. Baltic spring spawners show the lowest median and mean kinship coefficient. Baltic autumn show the second-lowest median kinship coefficient, but the mean is dragged higher by the high degree of relatedness in the Fehmarn population.

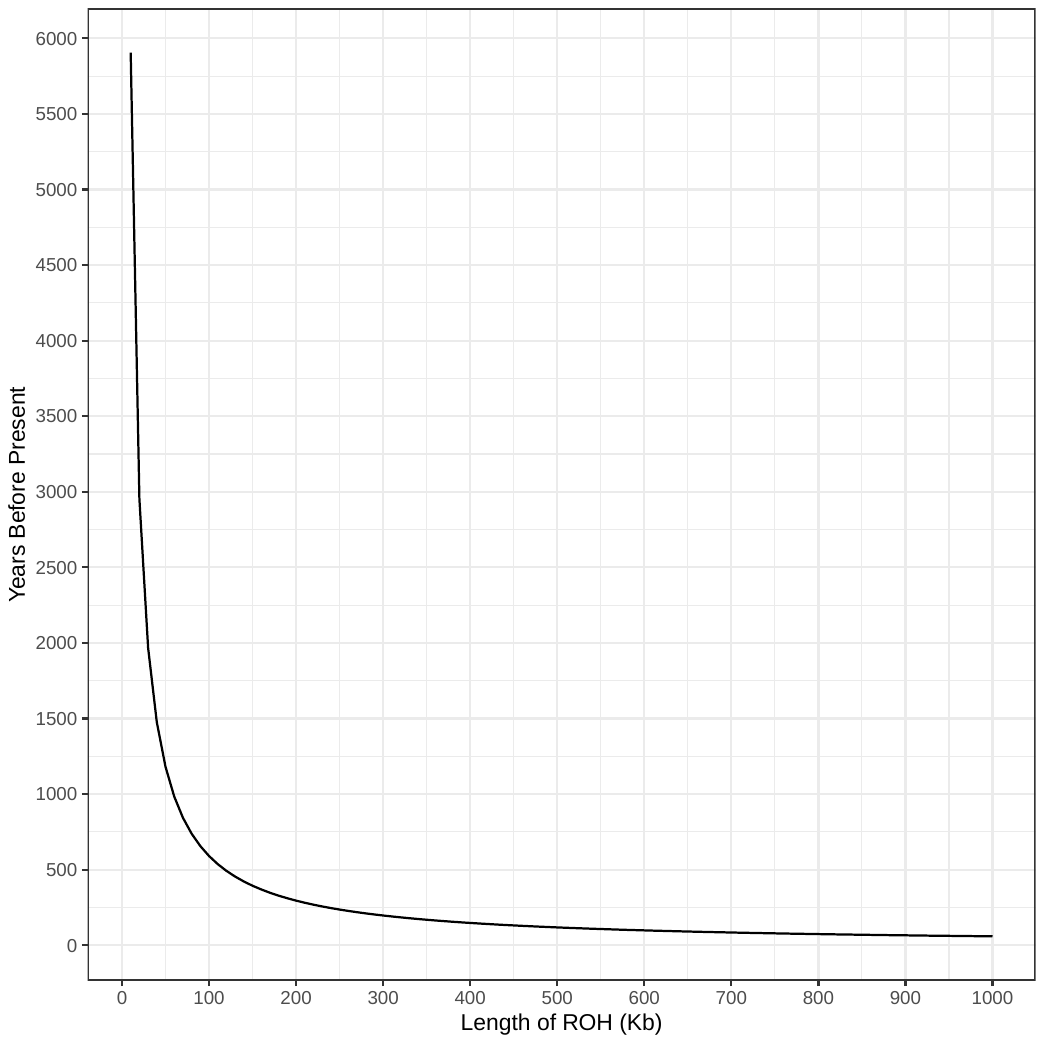

**Figure S11 – Time to coalescent event according to length of run of homozygosity.** Using the formula 100/2g cM/Mb = L^22^ and the known herring recombination rate of 2.54 cM/Mb, we calculated the relationship between length of ROH and timing of coalescent event. The curve indicates the length and its associated coalescent event in years before present.

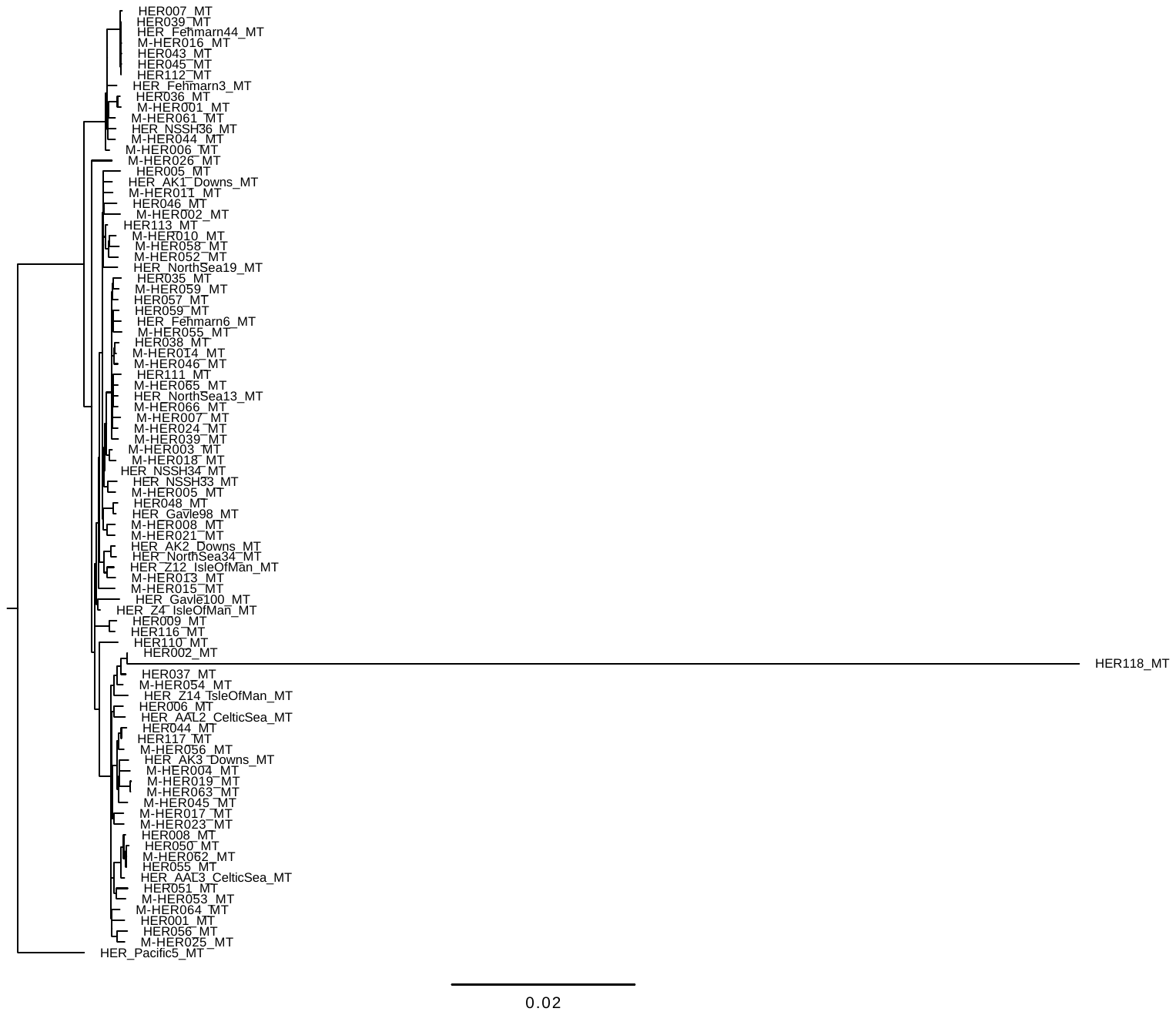

**Figure S12 – IQtree phylogeny of the herring mitogenome.** All ancient samples clustered with the modern samples and exhibited the structure reported by Teacher et al.^23^ that is not associated with geography. HER118 appears quite differentiated from the rest of the samples. BLAST showed that it is indeed a Baltic herring, therefore it was left in the analysis for BAMscorer assignment.

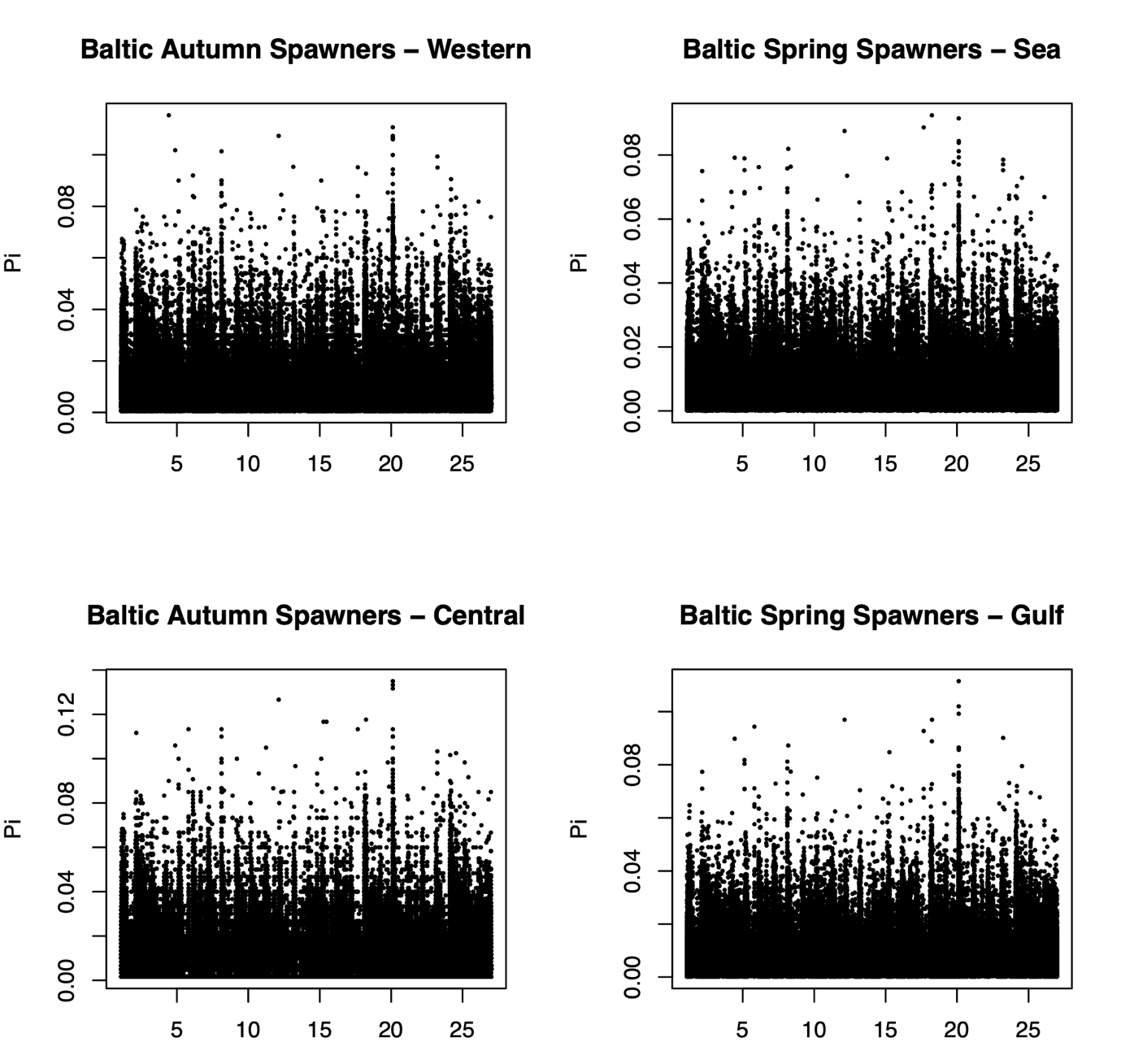
**Figure S13 – Estimates of** $\pi$ **for each of the four Baltic herring populations**. The western Baltic autumn spawners had an average pi of 0.006; the central Baltic autumn spawners had an average $\pi$ of 0.008; the Baltic spring spawners – sea had an average $\pi$ of 0.004; and the gulf spring spawners an average $\pi$ of 0.005.

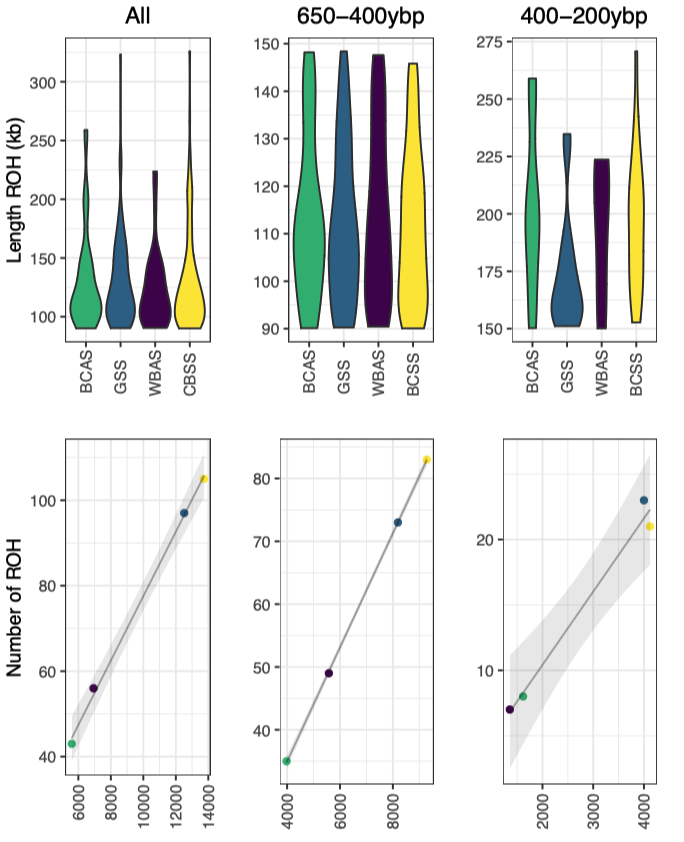

**Fig S14 – Runs of homozygosity in four modern Baltic herring stocks.** The top panel shows the distribution of ROH across the genome for the entire genome and two bins associated with coalescent events 650-400YBP and 400-200YBP. The bottom panel illustrates the summed ROH for each population compared to the total number of ROH. These results indicate a larger effective population size in the past for both autumn spawning herring populations and a smaller effective population size in the past for both spring spawning herring populations.

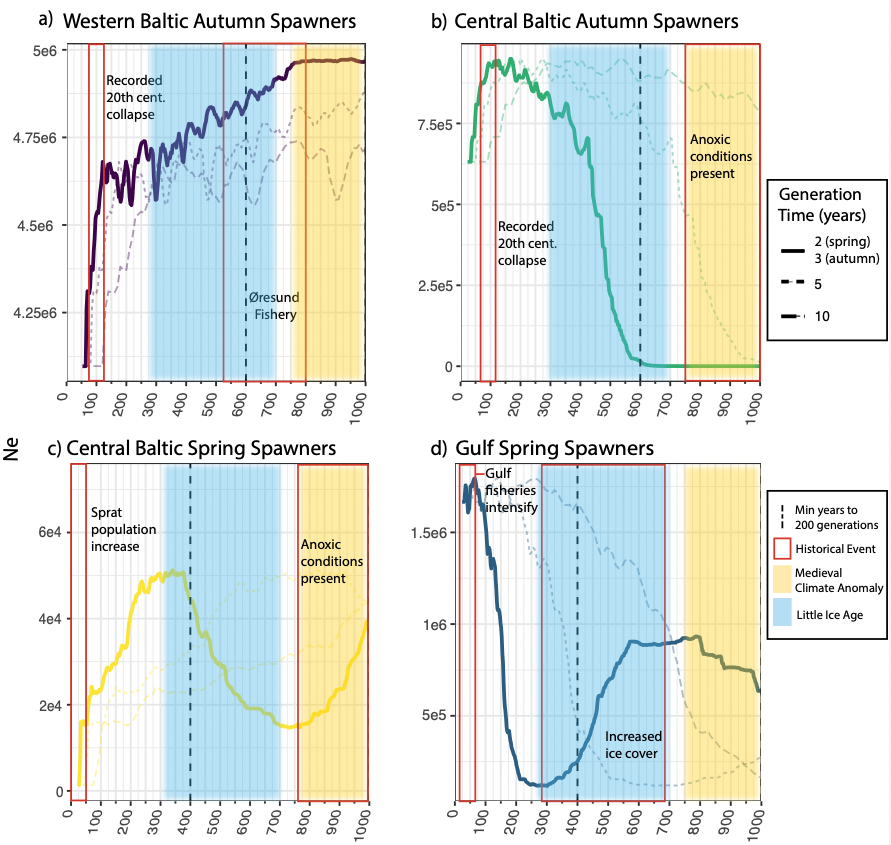

**Fig S15 – gone analysis with generation time uncertainty.** Each stock is represented in a separate chart: western Baltic autumn spawners (WBAS); central Baltic autumn spawners (CBAS); central Baltic spring spawners (CBSS); gulf spring spawners (GSS). Colored rectangles indicate key historical events. Yellow rectangles show the approximate duration of the Medieval Climate Anomaly (MCA) and blue rectangles the Little Ice Age (LIA). The dashed vertical lines show the minimum date (YBP) at which 200 generations in the past is reached (calculated using minimum generation times of 3 years for autumn spawners and 2 years for spring spawners), the known accurate window for *gone*. Historical events are denoted by red boxes for each population. As herring reproduce over their lifespan in overlapping generations, additional generation times were used to scale the demography for each population. These are visualized here as dashed lines. a) Demographic trajectory of the western autumn spawners shows a decline starting shortly after the start of the Øresund herring. They show an additional severe decline corresponding to the reported autumn spawning fishery collapse in the Baltic; b) Demographic trajectory of central autumn spawners, which appear limited during the MCA when anoxic conditions are present in the central Baltic. They increase during the LIA and rapidly decline during the period of known autumn spawning population collapse coinciding with the increase of the sprat (*Sprattus sprattus*) population; c) Central spring spawners show an increase around the time of the decline of the western autumn spawners, then a decrease again at the end of the LIA as well as another dramatic decrease around the time of the autumn spawners’ collapse ~100YBP; d) Gulf spring spawners decrease during the LIA and then increase dramatically at the end of the LIA, starting to decline only in very recent generations when fisheries in the gulfs intensify.

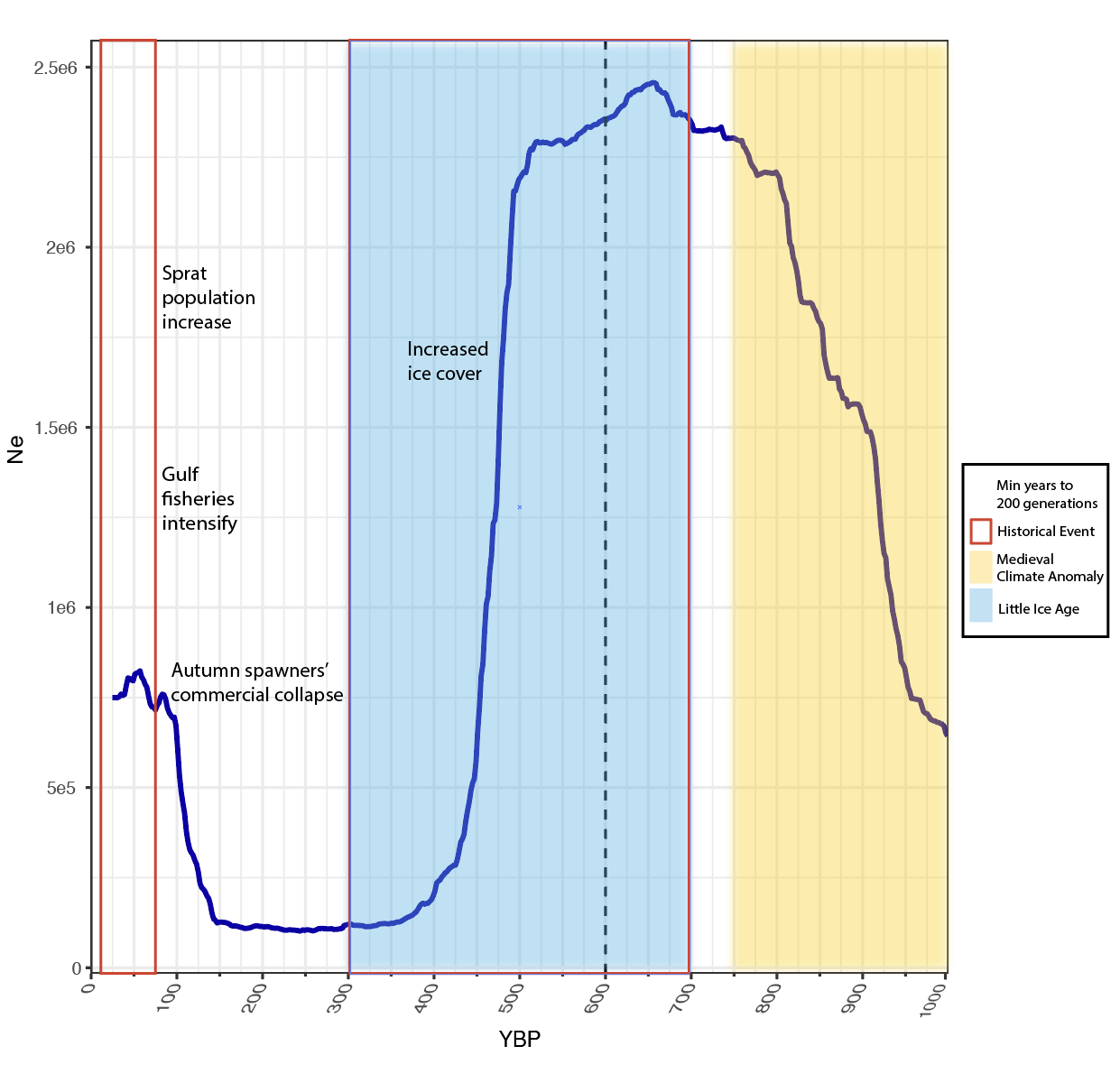

**Fig S16 – gone analysis for all Baltic spring spawners.** Here, Baltic spring spawners were grouped as a single metapopulation (n=10) and *gone* was run with the same parameters as before. This figure shows the projected demographic trajectory when both the GSS and CBSS are treated as a single population. Historical events are again marked, illustrating the relationship between fishing pressure, changing climate, and demographic trends in Baltic herring.
